# Coupling of cell proliferation to the segmentation clock ensures robust somite scaling

**DOI:** 10.1101/2025.01.10.632257

**Authors:** Marek J. van Oostrom, Yuting I. Li, Wilke H. M. Meijer, Tomas E. J. C. Noordzij, Charis Fountas, Erika Timmers, Jeroen Korving, Wouter M. Thomas, Benjamin D. Simons, Katharina F. Sonnen

**Author notes:** Division of Gene Regulation, the Netherlands Cancer Institute, Oncode Institute, Plesmanlaan 121, 1066CX Amsterdam, the Netherlands (Charis Fountas). Cell Biology, Neurobiology, and Biophysics, Department of Biology, Faculty of Science, Utrecht University, Padualaan, 3584 CS Utrecht, the Netherlands.

## Abstract

While scaling developmental processes is fundamental to maintaining robust tissue patterning, the mechanisms underlying this process are enigmatic. Somitogenesis, the periodic segmentation of growing mesodermal tissue in vertebrate embryos^1^, involves precise scaling with the unsegmented presomitic mesoderm (PSM) over developmental time and under perturbation^2–4^. Somitogenesis is spatiotemporally regulated by FGF and Wnt morphogen gradients and the segmentation clock — oscillations in Notch, Wnt, and FGF signalling^5–7^. Here, we find that cell proliferation is distributed throughout the oscillating PSM. Long-term single-cell tracking in mouse embryo tails uncovered a correlation between cell cycle progression and the segmentation clock, with microfluidics-based entrainment indicating coupling between the cell cycle and signalling oscillations, likely through S-phase inducing Cyclins. A theoretical model suggests this coupling ensures uniform PSM growth, uniform morphogen dilution and precise somite formation, which we validated experimentally by blocking cell proliferation. Our findings reveal that coupling cell proliferation to signalling oscillations is crucial for robust somitogenesis and precise somite scaling.

## Main

Somitogenesis is highly reproducible across individual animals of a given species^2^. Cells starting in the posterior tip of the embryo tail traverse the PSM bilaterally along the neural tube until they reach the anterior PSM, where differentiation and somite formation occur^1^. Imbalance between growth and segmentation can lead to premature or delayed arrest in somite formation, improperly sized somites or asymmetries between the left and right^3,4,8^, resulting in embryonic lethality or congenital disorders.

Scaling of somitogenesis, the relative adjustment of somite size to PSM size, occurs dynamically during mouse development: PSM size increases until embryonic day (E)9.5 and then decreases until somitogenesis terminates at (E)14.5^2^. At a given developmental time, variations in somite size are very low: at the 36-somite stage (E10.5) forming somites have a size of 162 ± 8 µm^2^, which corresponds to a maximum variation of only 1-2 cell diameters. Moreover, experimentally decreasing PSM size results in the formation of smaller somites^3,4^. How such accuracy and precision in segmentation of a variably sized tissue is achieved remains unclear.

The potential involvement of cell proliferation in somitogenesis has been a subject of discussion and controversy. While the cell cycle has been suggested to primarily contribute to PSM growth^9–11^, cell cycle perturbations do not interfere with segmentation clock oscillations in 2D *in vitro* culture models of human somitogenesis^12^. On the other hand, in zebrafish embryos, the absence of proliferation impairs proper somite formation and has been described as a source of noise for the segmentation clock^2,8,12–14^. A link between the segmentation clock and cell division has been suggested^13–15^, but not formally shown.

Here, we use methods including long-term cell tracking, microfluidics-based entrainment, and modelling to analyse how cell cycle dynamics impact somite scaling. Our data indicate that proliferation and signalling oscillations are coupled, and important for robust somite scaling in the context of somitic development.

## Results

### Cell cycle dynamics along the tailbud-to-somite axis

To understand the role of cell proliferation in somitogenesis, we first characterized cell cycle dynamics in uncultured mouse embryo tails. Immunostaining for the cell cycle marker Ki67 completely overlapped with DAPI staining, showing that all tailbud, PSM and somite cells were actively cycling (Fig. 1A). Next, we used flow cytometry to determine the distribution of single cells from uncultured mouse embryo tails over the four cell cycle phases: G1, S (when DNA replication occurs), G2 and M phase (when cell division occurs). We specifically analysed unsegmented mesoderm by gating for Tbx6-positive cells and found that a majority of cells were in S phase (60.4%), while fewer were either in G1 (22.7%) or G2/M phase (16.9%, Fig. 1B, Fig. S1). Previously, we found that mitosis occurs in the entire PSM^10^. To quantify mitosis distribution more precisely along the anteroposterior (AP) axis from tailbud to somite, we performed immunostaining using the mitotic cell marker pHH3 (phosphorylated histone H3) in mouse embryonic tails transversally sectioned from posterior to anterior (Fig. 1C-E, Fig. S2). We confirmed that cells divided within the tailbud, PSM and somite region of the tail (Fig. 1E). Furthermore, there was a slight increase in the percentage of pHH3-positive cells within the first two forming somite pairs compared to the PSM and tailbud.

**Figure 1.**
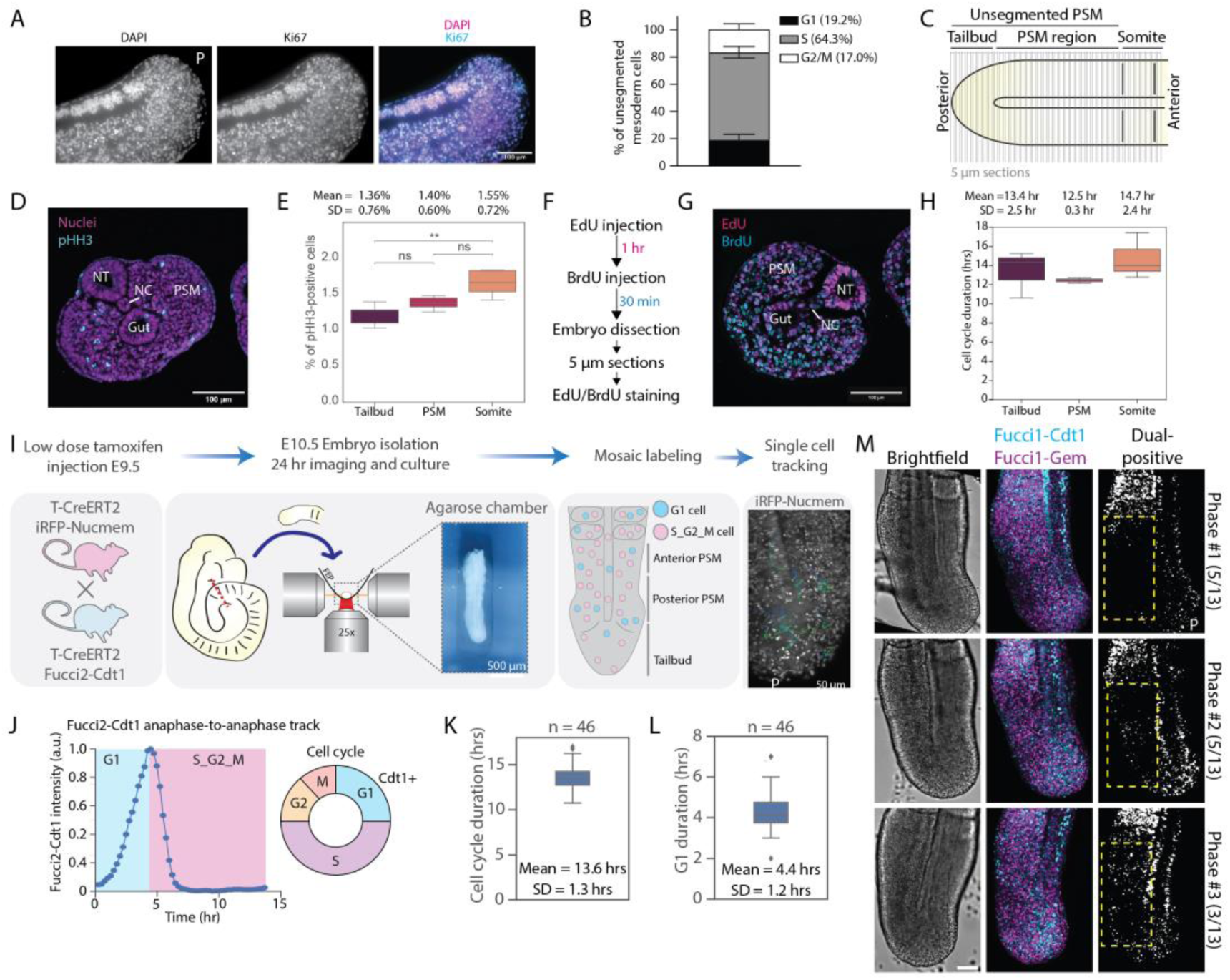
Cell cycle dynamics along the entire E10.5 presomitic mesoderm. **A-H** Quantification of cell cycle dynamics in uncultured tails. A 5 µm section of E10.5 tail with immunostaining for Ki67 and counterstaining with DAPI. **B** Flow cytometry analysis of embryonic tail cells stained for Tbx6 and DNA using Hoechst. N=3 experiments. Raw data shown in Fig. S4. **C-E** Quantification of late mitotic cells in embryonic tails. N=5. **C** Embryonic tails were sectioned into 5 µm sections. Immunostaining against phosphorylated-histone H3 (PHH3) and counterstaining with DAPI were performed. For the analysis, sections were divided into tailbud (posterior to neural tube), PSM and somite regions. **D** Representative image of a section stained against PHH3 and with DAPI (same section as in Fig. S2A). **E** Quantification of cells in late mitosis (pro-metaphase to anaphase) in the regions indicated in **B**. **F-H** Quantification of cell cycle dynamics by EdU-BrdU double-labelling. N=3. **F** Workflow outlining experimental steps. Embryos were dissected and tails sectioned akin to **B** before EdU/BrdU staining. **G** Representative image of a section stained for EdU, BrdU and nuclei (same image as in Fig. S3A). **H** Quantification of cell cycle length in the regions indicated in **B**. Coefficient of variation CV: CV(Tailbud) = 0.19; CV(PSM) = 0.02; CV(Somite) = 0.16. N = 3. **I-N** Quantification of cell cycle dynamics in *ex vivo* cultured tails. **I** Schematic illustrating single cell tracking pipeline. Pregnant mice were injected with low doses of Tamoxifen on E9.5 for sparse nuclear labelling of cells. Embryos were dissected after 24h of incubation. Embryonic tails were dissected and imaged with an inverted light-sheet microscope. Embryonic tails were placed in agarose grooves for microscopy. Mosaically labelled cells with nuclear marker iRFPnucmem were followed over time in 3D. **J** Representative timeseries data of the dynamics of the G1 marker Fucci2-Cdt1 in a single cell tracked from anaphase to anaphase. **K, L** Quantification of cell cycle length and duration that cells were positive for the G1 marker Fucci2-Cdt1. **M** Patterns of G1-S transition in uncultured E10.5 embryonic tails (same image as in Fig. S4F). Cells double-positive for Fucci1-Cdt1 and Fucci1-Gem were determined in PSM of embryonic tails. Tails were ordered into three phases according to their staining pattern: Phase 1: no double-positive cells in PSM; Phase 2: stripe of double-positive cells in PSM; Phase 3: double-positive cells distributed over PSM. Number in brackets indicates number of tails in which specific staining pattern was observed. Brightfield (*left panel*), merge of Fucci1-Cdt1 and Fucci1-Gem (*middle panel*) and cells double-positive for Fucci1-Cdt1 and Fucci1-Gem (*right panel*) depicted. N=13. Scale bar 100 µm. P indicates the posterior end of the embryo.

Next, we quantified the proliferation rate along the AP axis in embryo tails using pulse chase labelling by intraperitoneal injection of thymidine analogues 5-Ethynyl-2′-deoxyuridine (EdU) followed by 5-Bromo-2′-deoxyuridine (BrdU) 1 h later. This treatment was followed by imaging transversally sectioned tails^16,17^ (Fig. 1C, F-H, Fig. S3). The differentially labelled cell fractions allowed us to quantify cell cycle and S-phase duration per section (Fig. S3B, E). Cell cycle duration within the tailbud and PSM was approximately 13.5 hr (Fig. 1H, Fig. S3C) and S phase duration 10 hours (Fig. S3F-H). With a segmentation clock period of approx. 2.7 hours this indicates that it takes about 5 segmentation cycles for tissue size to double. Interestingly, variability in cell cycle duration was much lower in the PSM compared to tailbud and somites (Fig. 1H), hinting at tighter cell cycle control in the PSM.

Thus, cells divide along the whole AP axis at similar rates, suggesting constant expansion of the entire tailbud and PSM region. This contrasts with a model in which cells are added in the posterior PSM, while segmentation occurs in the anterior.

### Quantification of cell cycle dynamics using live-cell imaging

To study the dynamics of cell cycle progression at a single-cell rather than population level, we established a long-term single-cell tracking pipeline in the developing mouse embryo tail. To make this possible, we combined light-sheet microscopy with tissue stabilization in agarose grooves and mosaic labelling of cells (Fig. 1I, Fig. S4A-C). We crossed mice carrying the inducible Fucci2 cell cycle G1 reporter R26R-mCherry-hCdt1(30/120)^18^ (Fucci2-Cdt1) with mice carrying the inducible nuclear marker R26R-iRFPnucmem^19^ (iRFPnucmem) and a CreERT2 expression cassette driven by a T/Brachyury promotor (T-CreERT2)^20^. We then induced mosaic labelling by injecting low concentrations of Tamoxifen into pregnant female mice. Recombination resulted in approximately 10% of cells labelled with mCherry, which tracks the G1 phase of the cell cycle, and nuclear iRFP to allow tracking over time. Single cells were tracked within tissue explants and Fucci2-Cdt1 intensity was quantified over time (Fig. 1J, Movie S1). Cell-cycle duration was 13.6 ±1.3 hr, in agreement with the cell cycle duration *in vivo* (Fig. 1K), and cells spent on average 4.4 hr in G1 phase (Fig. 1L). Notably, we did not detect changes in cell density despite cell divisions over 24 hr of real-time imaging (Fig. S4D, E), indicating steady state tissue growth with cells continuously growing and dividing.

To determine whether cell cycle progression in the PSM is governed by spatiotemporal regulation, we analysed the spatial organization of cells double-positive for both the Fucci cell cycle reporters mKO2-hCdt1(30/120) (Fucci1-Cdt1) and mAG-hGem(1/110) (Fucci1-Gem) in mice expressing both transgenes in all cells. The presence of both Fucci1-Cdt1 and Fucci1-Gem signals in cells specifically marks the G1-S transition^21^. Notably, we identified 3 distinct phases for double positive cells in fixed tail explants: an absence of double positive cells in the PSM (Phase 1), a stripe of double positive cells in the PSM (Phase 2) and a wider spread of double positive cells throughout the PSM (Phase 3) (Fig. 1M, Fig. S4F). Such patterns are resonant with fixed images of cyclic segmentation clock genes showing different expression patterns in different embryos^22,23^. Our data therefore indicate that there might be a spatiotemporal organization of cell cycle entry in the PSM.

### The segmentation clock and cell cycle are correlated

To investigate whether there is a correlation between cell proliferation and segmentation clock oscillations, we simultaneously quantified cell cycle progression and segmentation clock oscillations in single cells using a mosaic labelling approach similar to the above. Specifically, we crossed mice carrying the inducible nuclear marker R26R-H2B-mCherry^24^, the T-CreERT2 transgene and the bright segmentation clock reporter Achilles-Hes7^25^, which is based on the Notch target gene and transcriptional repressor Hes7 (Fig. 2A-D, Fig. S5). We tracked single cells using the mosaically expressed mCherry signal (representing H2B), aligned intensity tracks to the timepoint of cell division and analysed Hes7 expression (Fig. 2C). The two daughter cells of a given mother remained closely related in terms of oscillation phase for up to 8 hours after division (Fig. S5C, Fig. 2A). Interestingly, we found that the oscillation phase of Hes7 one hour before division was not randomly distributed (timepoint 1, Fig. 2C, D). Rather, at the point of cell division, there was a sudden drop in nuclear Hes7 signal due to nuclear envelope breakdown. After division, individual cells immediately showed Hes7 oscillations again (Fig. S5A), but aligned tracks were not synchronized for approximately 4 hr (timepoint 2, Fig. 2C, D), after which cells started to synchronize again (Fig. 2C, D, Fig. S5E). The timeframe of non-synchronization after division roughly corresponds to the time cells spend in the G1 phase of the cell cycle (4.4 hr, Fig. 1L).

**Figure 2.**
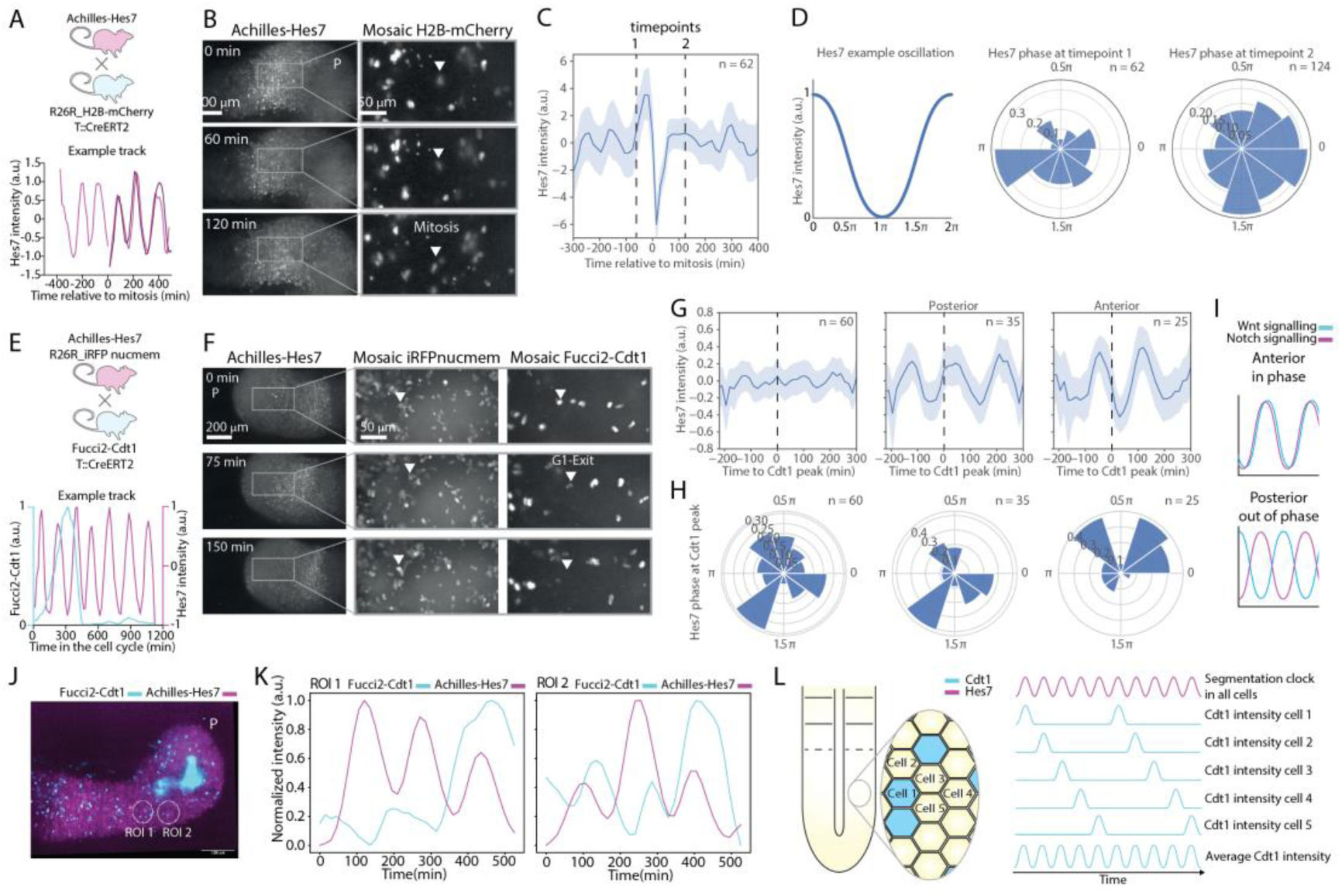
Cell cycle dynamics correlate with segmentation clock. **A-D** Correlation of Hes7 segmentation clock with cell division. **A** Experimental approach for sparse labelling of nuclei with H2B-mCherry combined with Achilles-Hes7. **B** Representative snapshots of timeseries data. Arrowhead points at one cell tracked over time. **C** Mean (blue line) and standard deviation (light blue shading) of Achilles-Hes7 signal of cells aligned to timepoint of cell division. **D** Schematic representation of a single phase wave. Phase of Achilles-Hes7 oscillations at timepoint 1, as indicated in **C**, (*left panel*) and at timepoint 2, as indicated in **C**, (*right panel*). **E-G** Correlation of Hes7 segmentation clock to G1-S transition. **E** Experimental approach for sparse labelling of nuclei with iRFPnucmem and Fucci2-Cdt1 combined with Achilles-Hes7. **F** Representative snapshots of timeseries data. Arrowhead points at one cell tracked over time. **G** Mean (blue line) and standard deviation (light blue shading) of Achilles-Hes7 signal of cells aligned to timepoint of G1-S transition. Left panel: all cells; middle panel: cells in posterior PSM; right panel: cells in anterior PSM. **H** Phase of Achilles-Hes7 oscillations at G1-S transition. Left panel: all cells; middle panel: cells in posterior PSM; right panel: cells in anterior PSM. **I** Schematic of the phase change in Wnt and Notch signalling between anterior and posterior PSM. **J** Maximum Intensity projection of E10.5 embryonic tail (purple: Achilles-Hes7, cyan: Fucci2-Cdt1). **K** Quantification of Achilles-Hes7 and Fucci2-Cdt1 in regions of interest (ROIs) as indicated in **J**. **L** Scheme illustrating correlation of cell cycle (duration 13.6 hr) to segmentation clock oscillations (2.7 hr).

The G1 restriction point is a point of no return, after which the cell cycle can run autonomously, even though later restriction points can prevent cell cycle progression to ensure the fidelity of cell division^26^. We therefore addressed whether segmentation clock oscillations were correlated with the G1-S transition. To this end, we combined Achilles-Hes7 with Fucci2-Cdt1 and iRFPnucmem and quantified expression in single cells of *ex vivo* cultured embryo tails after induction of mosaic labelling (Fig. 2E-G, Fig. S6A-D). We first analysed the period of segmentation clock oscillations along the posterior-anterior PSM axis and detected an increase in period towards the anterior (Fig. S7A), consistent with an oscillation period gradient^27^. We then separated cell tracks into their respective cell cycle phase based on Fucci2-Cdt1 reporter expression. When directly comparing all cells in G1 to all cells in the S/G2/M phase, independent of their location in the PSM, we detected a slightly higher period in S/G2/M phase cells (Fig. S7B). However, if we considered the position of the cells along the AP axis, we found that the period gradient was less pronounced in G1 than in S/G2/M phase cells (Fig. S7C,D). This indicates that segmentation clock oscillations change with both anteroposterior position in the PSM and cell cycle phase, suggesting an effect of cell proliferation on the segmentation clock^13,28^.

Next, we tested for a potential correlation between segmentation clock oscillations and the G1-S transition. When we aligned all tracks to the G1-S transition, defined as the peak in Fucci2-Cdt1 signal (Fig. 1J), we did not observe any relationship with the phase of Hes7 oscillations (Fig. 2G, Fig. S6E, F). In mice, the segmentation clock consists of oscillations in Notch, Wnt and FGF signalling. Previously, we demonstrated that the phase-relationship between Wnt and Notch signalling oscillations changes along the posterior-anterior axis of the PSM from out-of-phase in the posterior to in-phase in the anterior^29^ (Fig. 2I), corresponding to a 1π shift in phase-relationship. If an equivalent shift in phase-relationship exists between the cell cycle and Hes7 oscillations, any correlation would be masked when analysing the signal averaged along the anteroposterior axis. We therefore separated tracks from cells residing in the posterior and anterior PSM and analysed whether Fucci2-Cdt1 correlated with Hes7 dynamics. Notably, we found that the Fucci2-Cdt1 peak and the segmentation clock were correlated in both regions (Fig. 2G, Fig. S6E, F). In posterior PSM, the phase of Hes7 oscillations was ∼1.3π at G1-S transition, while it was ∼0.4π in anterior PSM. This corresponded to a change in phase-relationship between Notch oscillations and cell cycle of ∼0.9π between posterior and anterior PSM, roughly matching the 1π shift in Wnt-Notch phase-relationship. This provides indirect evidence that cell cycle progression correlates with Wnt-signalling oscillations in the PSM.

While the cell cycle in a single cell takes approximately 14 hr and a single cell spends approximately 4 hr in G1 phase, at the population level we observed an oscillatory dynamic in the Fucci2-Cdt1 reporter similar to the segmentation clock. Specifically, when quantifying small regions containing approximately 10-15 cells, we observed oscillations in Hes7 and Fucci2-Cdt1 signal with a period of roughly 2.5 hr (Fig. 2J,K, Movie S2). Such behaviour is consistent with a model in which groups of cells in this region correlate their G1-S transition in a manner that varies periodically between other neighbouring groups. This in turn implies that neighbouring cells within these regions were shifted in their G1-S transition by a multiple of complete segmentation clock periods relative to their neighbours (Fig. 2L).

Altogether these data reveal that cell cycle progression is correlated with signalling of the segmentation clock. More specifically, there is a constant phase-relationship with Wnt-signalling oscillations along the AP axis of the PSM.

### The segmentation clock and cell proliferation are coupled

We next sought to determine whether cell proliferation is merely correlated to the segmentation clock or whether there is a functional link between the two. Continuous inhibition of Wnt signalling by genetic or chemical perturbation leads to impaired PSM growth and a rapid regression of the PSM due to accelerated differentiation and loss of the morphogen gradient^30,31^ (Fig. S8). Such continuous perturbations are therefore not well suited to reveal a functional relationship between the segmentation clock and the cell cycle. Therefore, we aimed for a system to subtly modulate oscillations. To achieve this, we used our previously established microfluidic system to entrain oscillations to external pulses of pathway modulator^29,32^. We cultured embryonic tail tips in fibronectin-coated microfluidic chips, resulting in the spreading of the embryonic tissue to form a quasi 2D monolayer (Fig. 3A). In this system, the posterior-anterior embryonic axis is reorganized radially from centre to periphery^3,29^. Combining this set-up with mosaic labelling allows single-cell tracking and quantification of dim reporters, even by confocal microscopy^19^. Using embryos expressing Achilles-Hes7, inducible Fucci2-Cdt1 and iRFPnucmem^18,25,33^, we addressed whether a functional link exists between proliferation and the segmentation clock. Importantly, the correlation between segmentation clock (Hes7) oscillations and cell cycle (Cdt1) at the population level was maintained in the 2D cultures (Fig. S9, Movie S3).

**Figure 3.**
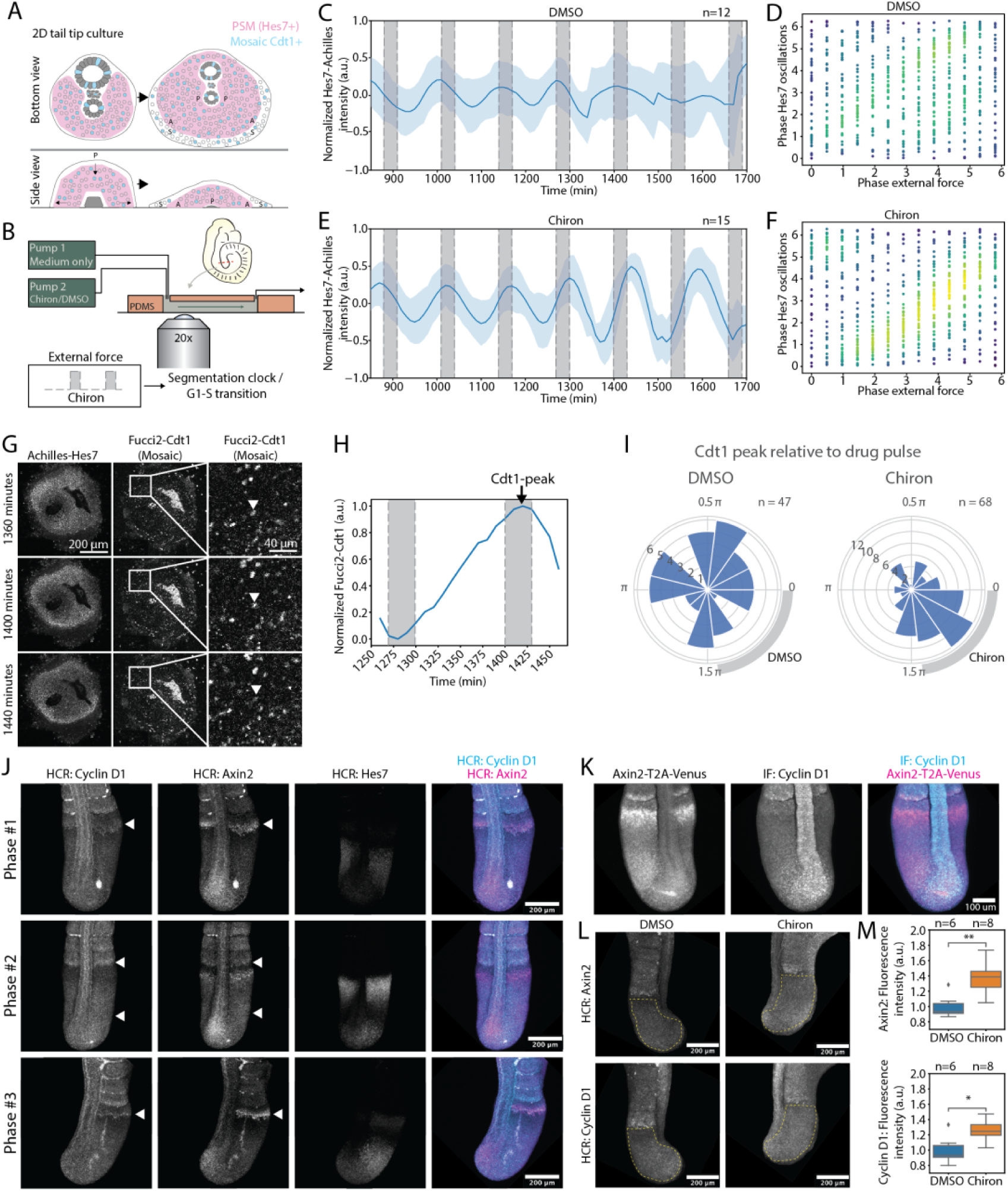
Coupling of cell proliferation to segmentation clock oscillations. **A-H** Microfluidics to test functional link. **A** Schematic illustrating a 2D tail tip culture spreading over time in the side view and bottom view of the embryo. **B** Experimental approach: Microfluidic system to modulate segmentation clock oscillations in *ex vivo* tailbud cultures. Analysis of segmentation clock and cell proliferation. **C-F** Analysis of segmentation clock: **C,E** Mean (blue line) and standard deviation (light blue shading) of Achilles-Hes7 signal in anterior PSM upon treatment with DMSO (**B**) and 5 µM Chiron (**E**). **D,F** Phase of Hes7 oscillations plotted against phase of external drug pulses for treatment with DMSO (**D**) and 5 µM Chiron (**F**). **G** Snapshots of representative fluorescence real-time imaging data highlighting tracking of a single cell (arrowhead). **H** Representative timeseries data showing Fucci2-Cdt1 reporter intensity of a single cell during microfluidic experiment. Grey shading indicates external drug pulse. **I** Phase of external drug pulses, at which G1-S transition occurs in DMSO-(*left panel*) or Chiron-treated samples (*right panel*). **J** Staining for mRNA levels of Axin2, Cyclin D1 and Hes7 in E10.5 embryonic tails using hybridization chain reaction (HCR) (same image as in Fig. S11A). **K** Immunostaining against Cyclin D1 in Axin2T2AVenus-expressing E10.5 embryonic tails (same as in Fig. S11B). **L-M** Treatment of E10.5 embryonic tails with DMSO or Chiron (10 µM). HCR for Axin2 (upper panels) or Cyclin D1 (lower panels). **L** representative images. **M** Quantification of Axin2 and Cyclin D1 in regions of interest as indicated in representative images. * < 0.05; ** < 0.01.

We then modulated the segmentation clock by periodically perturbing Wnt signalling using the GSK3 inhibitor Chiron (CHIR99021) (Fig. 3B, Movies S4). Our previous experiments had revealed that Chiron efficiently entrains Wnt-signalling oscillations, which also affects Notch-signalling oscillations of the segmentation clock^29^. Indeed, we confirmed that Chiron pulses entrained the segmentation clock using Achilles-Hes7 as a reporter (Fig. 3C-F). Subsequently, we analysed when the G1-S transition occurred (based on the peak Fucci2-Cdt1 reporter signal) within one signalling oscillation. We found that the G1-S transition preferentially coincided with drug pulses (Fig. 3G-I, Fig. S10). Interestingly, our previous experiments examining the dynamics of the Wnt-signalling reporter Axin2T2A revealed that this signal also peaked with Chiron pulses^29^, suggesting that the G1-S transition coincides with peaks in Wnt-signalling oscillations. In contrast, DMSO pulses led to a random distribution of Fucci2-Cdt1 peaks during the oscillation cycle (Fig 3I). Thus, modulating segmentation clock oscillations via targeted Wnt signalling modulation leads to an entrainment of cell cycle oscillations.

All signalling pathways of the segmentation clock are known mitogens promoting cell cycle entry. Recently, myc and Cyclin D, well-studied for their role in cell cycle progression and G1-S transition^34^, were identified as potential cycling genes at the mRNA level^6^. Indeed, using hybridization chain reaction (HCR), we found Cyclin D1 to be expressed in the PSM with a dynamic pattern overlapping with the Wnt-signalling target gene Axin2 (Fig. 3J, Fig. S11A). Since cyclins are post-translationally regulated by modulating protein stability, we stained for Cyclin D1 protein. The staining pattern for the protein also corresponded to the Axin2 reporter (Fig. 3K, Fig. S11B). To determine a functional link, we perturbed Wnt signalling and analysed the effect on Cyclin D1 expression. Since Wnt inhibition leads to an immediate loss of PSM due to differentiation, we instead activated Wnt signalling using Chiron. This led to an increase in both Axin2 and Cyclin D1 expression in the entire PSM within 5 hours of activation (Fig. 3L, M).

We conclude that the segmentation clock and cell proliferation are mechanistically coupled, presumably via periodic induction of cell cycle regulators, such as Cyclin D1, by Wnt-signalling oscillations.

### An extended somitogenesis model suggests that cell cycle-segmentation clock coupling ensures the fidelity of somite formation

To determine whether the coupling of cell proliferation to signalling oscillations might have a functional role in maintaining periodic somite formation, we turned to a conceptual modelling-based approach, extending on previous models of somitogenesis^4,35–37^. To develop the model, we placed emphasis on the following assumptions based on our findings and available literature: (1) Somite formation occurs at a certain phase between Wnt- and Notch-signalling oscillations and a low threshold of the FGF morphogen gradient^29,38,39^. (2) All PSM cells have a certain probability of dividing, and the cell cycle period takes values in the range 13.6 ± 1.3 hr with a refractory period of around 9.2 ± 0.5 hr after cell division, translating to the estimated length of the S/G2/M phase of the cell cycle (Fig. 1K, L). (3) Since FGF mRNA is known to be produced only in the posterior tailbud^39,40^, it follows that the FGF morphogen gradient is diluted by both mRNA degradation and cell division, during which mRNA is partitioned equally between the two daughter cells. (4) Only minimal rearrangement of cells in the one-dimensional model of the PSM was allowed^41^. This last assumption was consistent with quantifications of cell migration and rearrangement in the PSM, as tracked using light-sheet imaging (Fig. 4A-C, Fig. S12A-C): Cells in the tailbud moved constantly, with a slight preference in directionality towards the posterior end of the tissue (Movie S1, Fig. 4A). The velocity of cells moving away from the tail tip increased with increasing distance from the tail tip. In contrast, the movement of cells relative to their neighbours in the PSM was restricted (0.2 µm/ 15 min, Fig. 4B, C); i.e., cells changed their relative position by less than one cell diameter during a single segmentation clock cycle. Moreover, we did not detect a preferential division axis between the anteroposterior or mediolateral axis (Fig. S5D).

**Figure 4.**
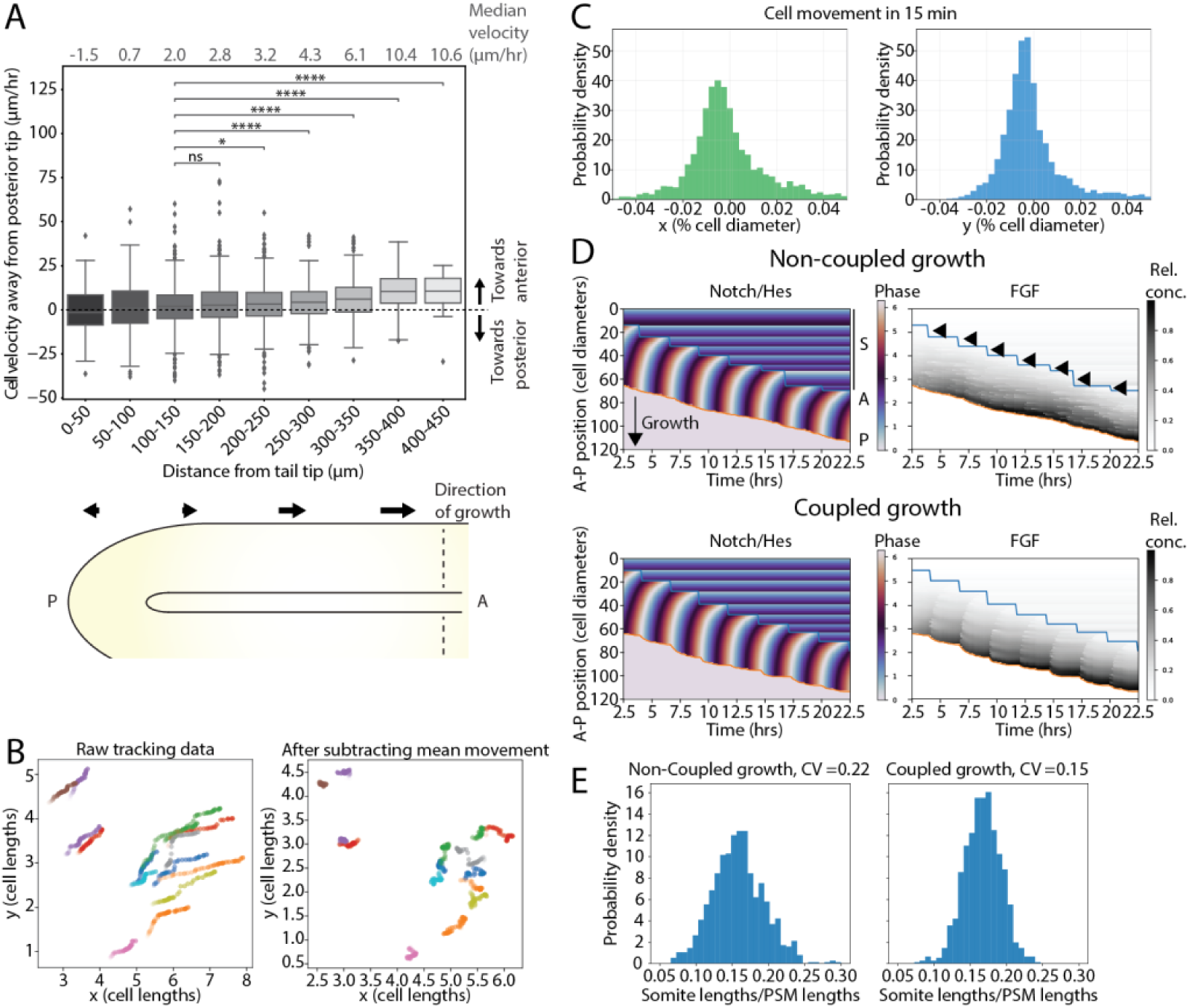
Coupling of cell proliferation to the segmentation clock promotes uniform PSM growth and somite formation in simulations. **A-C** E10.5 embryonic tails were cultured *ex vivo* and fluorescence real-time imaging was performed (corresponds to data from Figure 1,2). **A** Velocity of cells moving relative to the posterior tip (*top panel*). This indicates stretching of the PSM, as cells further away from the posterior have a higher velocity (*bottom panel*). Arrows indicate direction of movement. N = 6 tissues n = 106 tracks. * < 0.05; ** < 0.01; *** < 0.001; **** <0.0001. **B** Plots of single-cell tracks where whole-tissue movement was computationally removed to reveal movement of single cells relative to their surroundings. Colours correspond to different cells. Each dot represents location of a cell at specific timepoint (interval between timepoints 15 min). **C** Histogram of the amount of cell movement given in % of cell diameter (10 µm). N = 9 tissues n = 169 tracks. **D** Simulation of somitogenesis with non-coupled cell proliferation (top panels) or proliferation coupled to segmentation clock (bottom panels H). Example plots for simulations of signalling waves (*left panels*) as well as FGF gradient (gray shading) and somite formation (blue line) (*right panels*). Arrowheads indicate somite formation. Annotations are: Somites (S), Anterior (A) and Posterior (P). **E** Histogram showing scaling ratios, defined as the ratio of somite length to the PSM length based on 100 simulations (for details of the model and its parameters, see main text and Supplementary Theory). Coefficient of variation CV(non-coupled proliferation) = 3.5 cell diameters. CV(coupled proliferation) = 2.7 cell diameters.

Based on these results and assumptions, we modelled the dynamics of cells in the PSM as a one-dimensional line of Kuramoto-type phase oscillators coupled to local FGF density (for details of the model and its dynamics, see Supplementary Theory). Numerical simulations of the model recapitulated features of the continuously growing PSM field, showing the characteristic phase wave dynamic and periodic somite formation. Based on our observation that cell proliferation occurs along the entire PSM, we first asked what effect coupling between cell cycle entry and the segmentation clock has on somitogenesis and tissue patterning. To this end, we considered a line of 60 cells based on the number of cells in a line from posterior to anterior PSM and minimal cell exchange with neighbours (Fig. S12A-C and Fig. 4A-B). We then simulated tissue growth and somite formation (Fig. 4D top panels, Fig. S13). When cell divisions were allowed to occur randomly, decoupled from the segmentation clock, with a period of 13.6 ± 1.3 hr, numerical simulations revealed large variations in the somite scaling ratio, the ratio of the somite length over the PSM length (Fig. 4E, mean = 1.6, CV = 0.22). Moreover, uncoupled growth led to a non-uniform dilution of the FGF morphogen gradient (Fig. 4D, Fig. S13A). Conversely, when we imposed the condition that cell division occurs 1.25 hr after Wnt oscillations peak (5) (Fig 2D), tissue growth and somite formation were more robust, as evidenced by a sharper distribution of the somite scaling ratio (Fig. 4D, E, Fig. S13, B, mean = 1.7, CV = 0.15). This increase in robustness was observed consistently in simulations over a wide range of parameter values (Supplementary Theory). Together, these results suggest that the coupling of cell proliferation to the segmentation clock may be necessary to ensure robust tissue elongation and uniform dilution of the morphogen gradient, regulating the precise spacing of somite formation.

### Perturbing cell proliferation interferes with morphogen gradient dilution and scaling of somite formation

To understand how cell proliferation affects the segmentation process, we arrested cell proliferation in 2D *ex vivo* cultures^3^ of embryos expressing Fucci2-Cdt1, Achilles-Hes7 and iRFPnucmem using different cell cycle inhibitors. Inhibition resulted in an accumulation of non-dividing cells and an accumulation or absence of Fucci2-Cdt1, depending on when the arrest occurred during the cell cycle (Movie S5). Upon inhibition, segmentation clock oscillations were maintained (Fig. S14A, B), as shown previously^12^. This indicates that cell cycle progression is not a necessary driver of segmentation clock oscillations^12^.

Based on the minimal model, simulations suggest that in the absence of proliferation there would be a change in the scaling ratio from 0.171 in conditions of coupled growth to 0.138 in the absence of cell divisions (Fig. 5A). To challenge this prediction, we inhibited proliferation in growing embryo tails using a strong continuous S-phase inhibitor, Aphidicolin (Fig. 5, Movies S6). This led to a decrease in cell number, as expected, yet tissue size remained largely unaltered in the timeframe of the experiment due to continued cell growth, leading to a decrease in overall cell density (Fig. S14C-F). We then analysed the effect of Aphidicolin on somite formation. This treatment led to the formation of disorganized somites with undiscernible boundaries (Fig. 5B)^8^. In control cultures, the scaling ratio was 0.172 with a CV of 0.13, corresponding to the figure of CV = 0.15 for the model simulation. (Fig. 5C-E, 4E). Notably, consistent with the predictions of the model under perturbed conditions (Fig. 5A), the ratio changed from 0.172 to 0.109, indicating altered scaling of somite formation. We note that since (a) cell density differences between the PSM and somites (Fig. S14E-G) and (b) cell velocity (Fig. S14H) were the same under control conditions and following cell cycle inhibition, these cannot account for the change in scaling.

**Figure 5.**
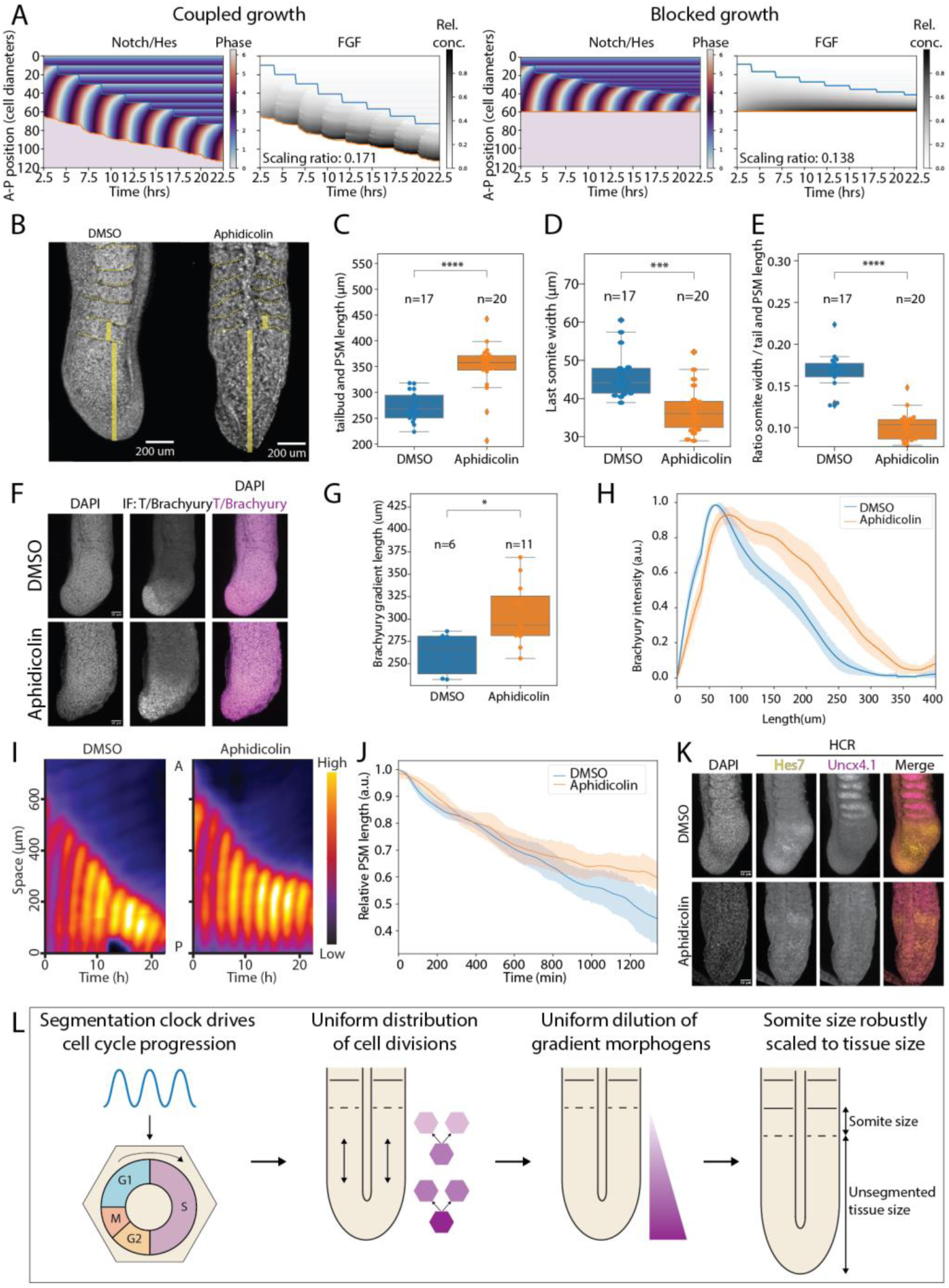
Coupling of cell cycle to segmentation clock affects scaling of somite formation. **A** Simulations of the somitogenesis model with cell cycle progression coupled to signalling (left) and following perturbation in which cell cycle is inhibited (right). Note the change in the average somite scaling ratio between the two conditions. **B-K** E10.5 embryonic tails were cultured *ex vivo* and treated with DMSO or the cell cycle inhibitor Aphidicolin. **B-E** After 23 hr of incubation, tails were fixed and stained with DAPI. **B** Representative images are shown. **C-D** Quantification of PSM length from tail tip to last formed somite boundary (**C**), somite length (**D**) and the ratio of somite length to PSM length as a measure of scaling (**E**) **F-H** After 23 hr of incubation, tails were fixed and stained for Brachyury and counterstained with DAPI. Scale bar 100 µm. **F** Representative images. **G** Quantification of the Brachyury signal along the AP axis of the PSM. Signal was normalized between 0 and 1. n(DMSO) = 6, n(Aphidicolin) = 11. **H** Quantification of the length of Brachyury gradient (Line: mean, shading: standard deviation). Intensity normalized between 0 and 1. * < 0.05; *** < 0.001; **** <0.0001. **I,J** Fluorescence real-time imaging of embryonic tails expressing Achilles-Hes7 was performed. **I** Representative kymographs of control and Aphidicolin-treated samples. **J** Quantification of PSM length by determining the size of the oscillating field (Line: mean, shading: standard deviation). Length normalized to the PSM length at experiment start. n(DMSO) = 9, n(Aphidicolin) = 11. **K** After 23 hr of incubation, tails were fixed and HCR against Hes7 and Uncx4.1 as well as counterstaining with DAPI were performed. Representative images are shown (same as in Fig. S14J). **L** Schematic model of how coupling of cell proliferation to segmentation clock affects somitogenesis. Morphogen density is indicated in magenta.

Finally, the model also predicted an expansion of morphogen gradients in the absence of cell divisions. We quantified morphogen gradients using T/Brachyury expression as a readout, given that this is directly affected by FGF and Wnt3a gradients^40,42^. Consistent with the model, the T/Brachyury gradient was expanded in the unsegmented PSM upon cell cycle inhibition (Fig. 5F-H). Similarly, the expression patterns of FGF8 and Axin2 mRNA were expanded and less graded than in controls (Fig. S14I-L). Consistent with the expansion of morphogen gradients^40^, the size of the oscillating field, a measure of the extent of differentiation, decreased less than in control tails (Fig. 5I-K, Fig. S14M). Furthermore, we found that the wave velocity did not decrease to the same extent as in control tissues but remained higher, despite the decrease in cell number relative to control tissue (Fig. S14M, N). This contrasted with unperturbed tissues, in which the wave velocity slows with decreasing tissue size and cell number^3^. While the density of cells decreased, the wave velocity increased in perturbed conditions. This implies that the relative phase-shift between oscillations in neighbouring cells decreased in the absence of cell division. Thus, absence of cell division impairs somite formation and scaling by affecting tissue gradients.

## Discussion

Here, we examined the role of cell proliferation in somitogenesis, uncovering a tight correlation between cell cycle entry and the phase of the segmentation clock oscillation. Further, we hypothesized and validated a mechanism by which the coupling of cell division to the segmentation clock may ensure uniform PSM growth and dilution of regulatory molecules, thereby modulating the scaling and accuracy of somite formation.

In contrast to studies that support a posterior growth zone^43–45^, we found that cell proliferation was uniformly spread over the posterior and anterior PSM, consistent with previous analyses^2,10^. Furthermore, cell proliferation was coupled to the segmentation clock, likely via Wnt-induced expression of pro-proliferative factors such as Cyclin D^6^, which ensures uniform distribution of proliferating cells over the PSM. With a proliferation rate of approximately 14 hr compared to a somitogenesis period of 2.7 hr, we found that cell proliferation is a key component of mouse somitogenesis. Upon cell cycle inhibition, cell division was abolished while tissue and cell growth continued, indicating that cell division in the PSM has a negligible effect on tissue growth within the timescale of the experiment.

Morphogen gradients are generated by mRNA and protein production in the posterior tip and levels slowly decrease as cells move down the posterior-anterior axis of the PSM^39^. Thus, morphogen gradients are modified by mRNA (and protein) degradation as well as cell division, which leads to a continuous decrease in morphogen concentration and nuclear effector proteins in cells. Indeed, inhibiting cell division in the tissue expanded the morphogen gradients. In agreement with the clock-and-wavefront model^37^, regression of the oscillating field was also reduced upon cell cycle inhibition. Moreover, signalling wave velocity was increased and somite formation impaired. This suggests that the principal role of cell division in the PSM may not lie in its contribution to tissue growth but rather in shaping morphogen gradients for proper somite formation.

This is consistent with a model (Fig. 5L) in which cell divisions are uniformly distributed over the tissue by coupling to the segmentation clock. This leads to an increase in cell number uniformly distributed over the PSM, which leads to (1) a uniform expansion of the oscillating field and (2) a uniform dilution of morphogen mRNA and protein as well as downstream gradients. These gradients are known to affect wave dynamics^46^ and both in turn regulate somite formation^38,40^. Supported by mathematical modelling, this suggests that the coupling of cell divisions to the segmentation clock ensures the formation of consistently scaled somites. Scaling of segment size to PSM size has previously been correlated with scaling of wave dynamics^3^ or the FGF gradient^4^ in mouse and zebrafish, respectively, both of which we found to be affected by cell cycle inhibition. Thus, this constitutes a mechanism by which local cell-intrinsic regulations, i.e., whether to divide or not, impact the tissue-wide regulation of morphogenesis to ensure robust pattern scaling and somite formation.

How our findings translate to other species remains to be addressed. In zebrafish, continued tissue growth, impaired somite formation and decreased cell density are observed in the PSM of *emi1* mutants, which stop proliferation during gastrulation^8^. Moreover, segmentation clock oscillations correlate with cell divisions in zebrafish^14^, even though cell proliferation is thought to play a minor role for tissue growth^47^. The coupling we observed could explain why zebrafish segmentation proceeds more quickly than growth and oscillation regression, leading to a rapid reduction of tissue size and a decrease in oscillation wave amplitude from 4π to 2π over developmental time^36^. In human somitogenesis, a correlation between the timepoint of cell division and Hes7 oscillations has so far not been observed^12^. To clarify the role of cell proliferation in human cells, cell cycle reporters specific for the G1-S transition should be quantified relative to Hes7 oscillations in posterior vs. anterior PSM cells.

Signalling oscillations have been observed in other tissue types and developmental contexts. Correlations between such oscillations and cell division have also been described^48^. For instance, in *in vitro* cultures of the small intestine, we recently identified Hes1 oscillations that correlated with cell divisions^49^. In neural stem cells, interactions between Hes1 oscillations and the cell cycle inhibitor p21 have been described^50^. Whether these other systems depend on active coupling between the cell cycle and signalling oscillations, as shown here for somitogenesis, and whether this coupling impacts growth, differentiation and morphogenesis at the tissue level are exciting questions for future research.

## Methods

### Animals and Housing

All experimental procedures were conducted in accordance with institutional guidelines and approved by the Ethical Review Board of the Hubrecht Institute. Animal protocols were reviewed and authorized by the Animal Experimentation Committee (DEC) of the Royal Netherlands Academy of Arts and Sciences (KNAW). Mice were bred and housed under standard conditions at the Hubrecht animal facility, with a controlled environment maintained at 50–60% humidity, 22–23°C temperature, and a 12-hour light/dark cycle. Mice had ad libitum access to water and standard chow throughout the study.

### Mouse Strains

All transgenic mouse lines used in this study have been previously described. The Achilles-Hes7 line was generously provided by the Kageyama laboratory^25^. Fucci ^21^, The R26R-H2B-mCherry, and R26R-mCherry-hCdt1(30/120)^24^ lines were obtained from the RIKEN Center for Biosystems Dynamics Research and the RIKEN BioResource Research Center (accession numbers CDB0204K, RBRC02706, and CDB0229K, respectively) via the RIKEN Mouse Repository. The Tg(T-cre/ERT2)1Lwd/J (“T-CreERT2”) line was sourced from The Jackson Laboratory (accession number #025520). The R26R-iRFPnucmem^19^ and Axin-T2A-Venus^29^ lines were provided by the Aulehla laboratory. All wild-type mice used in this study were F1 hybrids (BALB/cJ × C57BL/6J).

### Animal and Embryo Work EdU/BrdU Incorporation

Female mice were sacrificed at 10.5 days post-coitum (dpc) and embryos were subsequently dissected. For EdU and BrdU incorporation experiments, pregnant females at 10.5 dpc were intraperitoneally injected with 3.3 mg EdU in PBS, followed one hour later by an equimolar injection of 4.0 mg BrdU in PBS^51^. Mice were sacrificed 30 minutes after BrdU injection. Embryos were dissected in PBS supplemented with 1% Bovine Serum Albumin (Biowest, P6154) for tail extraction.

### Mosaic Labelling

For mosaic labelling experiments, pregnant females were intraperitoneally injected with tamoxifen (Sigma-Aldrich, T5648, dissolved in corn oil) on embryonic day E9.5. To label with H2B-mCherry, 1 mg of tamoxifen was administered, whereas for labelling with iRFPnucmem and Fucci reporters, 0.1 mg of tamoxifen was used.

### Flow Cytometry

For flow cytometry analysis, E10.5 mouse tails were extracted and cut into smaller pieces using a surgical scalpel to facilitate single-cell dissociation. Tail fragments were washed three times in PBS0 in Eppendorf tubes, followed by a 15-minute incubation with 400 µl of TrypLE (Thermo Fisher, 12604013). After incubation, the tissue was passed through a 30G needle to generate a single-cell suspension. The suspension was then washed twice in PBS0 supplemented with 1% Bovine Serum Albumin (Biowest, P6154) by centrifugation at 500 g for 5 minutes.

To fix the cells, the pellet was resuspended in 4% paraformaldehyde (PFA) and incubated at room temperature for 15 minutes. After two additional washes in PBS with 1% BSA, cells were stored at 4°C until further staining. For immunofluorescent staining, the single-cell pellet was resuspended in 3% BSA PBS with 0.25% Triton X-100 and incubated overnight at 4°C with anti-Tbx6 rabbit primary antibody (Abcam, ab38883) at a dilution of 1:1000. After three washes in 3% BSA PBS, the cells were incubated with secondary donkey anti-rabbit Alexa Fluor 488 antibody (Thermo Fisher, A-21206) at a dilution of 1:1000 for 2 hours at room temperature.

Following a further two washes, the single-cell suspension was resuspended in 500 µl of PBS supplemented with Hoechst 34580 (1 µg/ml) and passed through a 20 µm mesh. Flow cytometry was performed on a CytoFLEX S Flow Cytometer, and the Tbx6-positive cell population was selected for cell cycle analysis using FlowJo software.

### Culture of Live Embryonic Tissue

Culture of whole mount and spread-out tail tip tissues was performed as previously described^29^. For perturbation experiments on whole mount and spread-out tail tips, the following compounds were used: CHIR99021 (Chiron, Sigma-Aldrich, SML1046) at 10 µM, IWP-2 (Sigma-Aldrich, I0536) at 1 µM, Aphidicolin (ThermoFisher, J60236.MCR) at 1.6 µg/mL, Thymidine (Sigma-Aldrich, T9250) at 2 mM, Palbociclib (Selleckchem, PD-0332991) at 1 µM, and RO-3306 (Sigma-Aldrich, 217699) at 10 µM.

### Microfluidics Experiment

The full protocol for performing the microfluidics experiment has been described elsewhere^29,32^. For entrainment of Achilles-Hes7 and Fucci2-Cdt1 reporter lines, a pulsing scheme was employed consisting of 100 minutes of medium followed by 30 minutes of medium containing CHIR99021 (Chiron, Sigma-Aldrich, SML1046) at 5 µM or DMSO.

### Confocal Live Imaging

Confocal live imaging was performed using a Leica TCS SP8 confocal microscope equipped with a HC PL APO CS2 20×/0.75 DRY objective. For cell tracking experiments, images were acquired every 10 minutes with a z-step of 5 µm, a stack depth of 50 µm, and a z-resolution of 4 µm. For non-tracking imaging, images were taken every 10 minutes with a z-step of 10 µm, a stack depth of 100–150 µm, and a z-resolution of 9 µm. Excitation of Achilles, mCherry, or iRFP signals was performed using OPSL lasers at 514 nm, 568 nm, or 638 nm, respectively. Movies were recorded at a resolution of 512 × 512 pixels (pixel size 1.517 µm). For wholemount tissues, imaging was conducted in a humidified atmosphere maintained at 37°C, with 5% CO₂ and 60% O₂. For 2D cultures the O₂ was reduced to 20%.

### Light-sheet Live Imaging

Live imaging of whole mount embryo tails for single-cell analysis was performed using a Viventis light sheet microscope (Viventis Microscopy, #LS1 Live). Imaging was conducted in a humidified atmosphere maintained at 37°C, with 5% CO₂ and 60% O₂. For fluorescence excitation, lasers with wavelengths of 514 nm, 568 nm, and 687 nm were used for Achilles, mCherry, and iRFP, respectively. Imaging was carried out with a 25× objective, using a laser beam of 3.3 μm and acquiring a z-stack of 60 steps, each with a 3 μm increment. Images were captured at a resolution of 2048 × 2048 pixels every 15 minutes. To prevent whole mount tissues from moving out of the field of view while still allowing unperturbed growth, a 2% agarose gel in PBS was cast around a custom-made insert in the sample holder. This insert created five spaces (1.5 mm × 0.5 mm) for the individual tissues to be loaded.

### Paraffin Sectioning of Tissues and Immunofluorescence

Tissues fixed overnight in 4% paraformaldehyde (PFA) at 4°C were stored in 70% ethanol at 4°C for up to one week prior to embedding. Embryo tails were then subjected to Xylene treatment, embedded in paraffin in groups of three, and sectioned transversally into 5 μm thick slices.

For staining, the slides were sequentially immersed in Xylene, followed by 100%, 95%, 70%, and 50% ethanol, and rinsed with running cold tap water for 3 minutes each. Antigen retrieval was performed by heating the slides in a citrate buffer (8.2 mM sodium citrate and 1.8 mM citric acid, pH 6). Slides were blocked and permeabilized using 1% Bovine Serum Albumin (BSA, Biowest, P6154) in 0.5% Triton X-100 PBS. Mitotic cells were stained with anti-phospho-histone H3 (Sigma-Aldrich, 05-806) at a 1:1000 dilution in 1% BSA PBS, and incubation was carried out overnight at 4°C. Secondary staining was performed using Donkey anti-Mouse Alexa Fluor 488 (Thermo Fisher, A-21202) at a 1:1000 dilution, supplemented with DAPI (5 μg/mL, Sigma) in 1% BSA PBS, and incubation was carried out overnight at 4°C. After staining, slides were mounted with Vectashield (H1200NB, Vector Laboratories). Images were acquired using a Leica TCS SP8 confocal microscope with an HC PL APO CS2 20×/0.75 DRY objective and OPSL lasers at 365 nm and 488 nm for excitation.

For BrdU and EdU staining, two additional steps were incorporated. Prior to blocking and after antigen retrieval, slides were incubated at room temperature in 2 M HCl for 10 minutes. EdU was then visualized using the Alexa Fluor™ 647 dye from the Click-iT™ EdU Cell Proliferation Kit for Imaging (Thermo Fisher, C10340). BrdU was detected using an anti-BrdU antibody (Thermo Fisher, B35128) at a 1:500 dilution, followed by secondary staining with Donkey anti-Mouse Alexa Fluor 568 (Thermo Fisher, A10037).

### Whole Mount Tissue Antibody Staining

Prior to immunostaining, samples were fixed overnight in 4% formaldehyde at 4°C and stored in PBS at 4°C. Primary antibodies used for staining included anti-Ki67 (Abcam, ab16667) at 1:1000, anti-T (human/mouse) (R&D Systems, AF2085-SP) at 1:1000, and anti-Cyclin D1 (Cell Signaling, 2978) at 1:500. Secondary antibodies included Donkey anti-Goat Alexa Fluor 488 (Thermo Fisher, A-11055) at 1:1000 and Donkey anti-Rabbit Alexa Fluor 568 (Thermo Fisher, A10037) at 1:1000.

Immunostaining was carried out by incubating the samples with primary and secondary antibodies overnight in PBST (0.1% Triton X-100) containing 1% Bovine Serum Albumin (BSA, Biowest, P6154). The secondary antibody mixture was supplemented with DAPI (Sigma, 5 μg/mL) for nuclear staining. For imaging, a z-stack consisting of 10-12 planes with a 10 µm distance between each plane was acquired at a resolution of 1024 × 1024 pixels (pixel size 0.758 µm) using a Leica TCS SP8 MP confocal microscope with a HC PL APO CS2 20×/0.75 DRY objective. Excitation was performed with OPSL lasers at 365 nm, 488 nm, and 568 nm.

### Whole Mount Tissue Hybridization Chain Reaction (HCR)

Hybridization chain reaction (HCR) was performed as described previously^52^. Cyclin D1, Hes7, Uncx4.1 and Fgf8 probe sets were generated using a probe generator (https://github.com/rwnull/insitu_probe_generator). Probe sets for Axin2, Hes7, and Uncx4.1 were obtained from Molecular Instruments. HCR imaging was performed using a Leica TCS SP8 confocal microscope with a HC PL APO CS2 20×/0.75 DRY objective. The system was equipped with OPSL lasers for excitation at 365 nm, 488 nm, 568 nm, and 638 nm. Probe sets for Axin2 were obtained from Molecular Instruments. HCR imaging was performed using a Leica TCS SP8 confocal microscope with a HC PL APO CS2 20×/0.75 DRY objective. The system was equipped with OPSL lasers for excitation at 365 nm, 488 nm, 568 nm, and 638 nm.

### Image and Data Processing

To quantify the fraction of phospho-histone H3 (pHH3)-positive cells, skin, neural tube, notochord, and gut were manually removed from the DAPI channel in section images using Fiji. Subsequently, StarDist, a Fiji plugin, was used to segment the remaining nuclei, providing an accurate measurement of the number of nuclei per slice. pHH3-positive cells were manually counted per slice.

For the quantification of EdU- and BrdU-positive cells, skin, neural tube, notochord, and gut were similarly excluded from the DAPI channel in Fiji. This channel was further denoised using Remove Outliers in Fiji (pixel size= 4, threshold=50) followed by a gaussian blur with a sigma of 2. Using Cellpose cryo3 model we segmented the remaining nuclei, allowing for an accurate count of the total number of nuclei per slice. Mean intensities of EdU and BrdU were calculated within the DAPI masks, and thresholds were applied to categorize cells as positive or negative for EdU and/or BrdU. These annotations, expressed as fractions of the total cell count, were used for the quantification of cell cycle duration, population doubling time, and S-phase duration (see Fig. 1, Fig. S2,3).

To visualize tissue-wide dynamics of Hes7 and Fucci2-Cdt1, regions of interest (ROIs) were drawn in maximum intensity projections of time-lapse movies in Fiji. Intensity measurements of both channels were quantified over time, smoothed, and linearly normalized between 0 and 1. Kymographs of tissue-wide dynamics were generated using the KymoResliceWide plugin in Fiji. A line was drawn from the posterior to the anterior end of the tissue, with a width of 100 μm.

For the quantification of the T/Brachyury gradient, a similar approach was used. A line plot was generated by measuring the intensity of T/Brachyury along a line drawn in Fiji. Gradient length was defined as the distance from the tip of the tail to the point where T/Brachyury intensity reached 10% above background level. To quantify staining intensities, an ROI was manually drawn within the PSM, and the mean intensity was calculated using Fiji. For the measurement of PSM and somite lengths, DAPI images were used. Lines were drawn parallel to the neural tube from the tail tip to the first somite boundary for PSM size, or from the first to the second somite boundary for somite size.

For the quantification of tissue density, an ROI was drawn within confocal or light sheet images, and the number of cells within the selected area was counted using either the Achilles-Hes7 or DAPI channel.

To track single cells over short time intervals in confocal microscopy movies, an ROI of 3 × 3 pixels was manually tracked within the Fucci2-Cdt1 or Achilles-Hes7 channel in Fiji. The peak intensity of Fucci2-Cdt1 was defined as the moment when a cell transitions from G1 to the S-G2-M phase. These tracks were used to quantify cell velocity or the timing of cell cycle transitions.

For long-term single-cell tracking in light sheet images, cells were manually tracked by following mosaically expressed H2B or Nucmem reporters using the Fiji plugin Mastodon (https://github.com/mastodon-sc/mastodon). Spot centre intensities were measured within the tracked spots to quantify Achilles-Hes7 or Fucci2-Cdt1. These intensities were smoothed using a moving average of 3 time points and linearly normalized, such that each track contained intensity values between 0 and 1. For the analysis of Hes7 dynamics, PyBOAT was used with the following settings: min_period = 60 min, max_period = 300 min, and TC = 150. All data plotting was performed using Python. For binning the cell positions along the tailbud-to-somite axis we tracked the posterior tail tip and the forming somite boundaries over time and quantified a relative AP position of the cells were at any given time. We considered 0.0-0.6 as posterior and 0.6-1.0 as anterior.

### Statistical Analysis

All experiments were conducted with a minimum of three independent replicates. Data visualization was performed using Tukey-style boxplots, where the bar represents the median, the box delineates the 25th and 75th percentiles, and the whiskers extend to 1.5 times the interquartile range (IQR). Violin plots were used to display the distribution of data from the maximum to the minimum, with the bar indicating the median value. For statistical comparisons between groups, normality was assessed using the Shapiro-Wilk test. Statistical significance was determined using either an independent t-test or the Mann-Whitney U test, depending on whether the data followed a normal distribution. The coefficient of variation CV is calculated from the standard deviation std and mean as follows: CV = std/mean.

## Data and Code availability

All data and custom code used in this study are available upon request to.

## Supporting information

Supplementary Theory

## Acknowledgements

We thank members of the Sonnen group for discussions and feedback. We are grateful to Takashi Hiiragi and Juan Garaycoechea for discussions and feedback on the manuscript. We thank Cantas Alev for discussions. We thank Jan-Daniël de Leede for help with data analysis. We thank the animal, FACS and imaging facilities of the Hubrecht Institute for their support. We thank Alexander Aulehla for support and providing mouse lines, in particular the inducible iRFPnucmem line. We thank Andrea Boni for help with setting up the embryo-tail imaging in the light-sheet microscope. The H2B-mCherry mouse line was kindly provided by the Riken Center for Biosystems Dynamics Research and the Fucci line by the Riken BioResource Research Center. The T-Cre/ERT2 line was kindly provided by Jackson Laboratories. The Achilles-Hes7 was kindly provided by Ryoichiro Kageyama. This work was supported by the Hubrecht Institute and received funding from the European Research Council under an ERC starting grant agreement no. 850554 to K.F.S. In addition, this research was supported in part by grant NSF PHY-1748958 to the Kavli Institute for Theoretical Physics (KITP). B.D.S. is supported by the Wellcome (219478/Z/19/Z) and the Royal Society (RP\R\231004).

## Author Contributions

M.J.O. and K.F.S. conceived the project together with Y.I.L. and B.D.S.; K.F.S. supervised the project; M.J.O. performed experiments with help from W.H.M.M., T.N., C.F., E.T., J.K. and W.M.T.; M.J.O., Y.I.L. and K.F.S. analysed the data; Y.I.L. and B.D.S. developed the theoretical framework; M.J.O., Y.I.L. and K.F.S. wrote the manuscript with input from all authors.

## References

1 Meijer, W. H. M. & Sonnen, K. F. From signalling oscillations to somite formation. Current Opinion in Systems Biology 39, 100520 (2024). 10.1016/j.coisb.2024.100520

2 Tam, P. P. The control of somitogenesis in mouse embryos. J Embryol Exp Morphol 65 Suppl, 103–128 (1981).

3 Lauschke, V. M., Tsiairis, C. D., Francois, P. & Aulehla, A. Scaling of embryonic patterning based on phase-gradient encoding. Nature 493, 101–105 (2013). 10.1038/nature11804

4 Ishimatsu, K. et al. Size-reduced embryos reveal a gradient scaling-based mechanism for zebrafish somite formation. Development 145 (2018). 10.1242/dev.161257

5 Hubaud, A. & Pourquie, O. Signalling dynamics in vertebrate segmentation. Nat Rev Mol Cell Biol 15, 709–721 (2014). 10.1038/nrm3891

6 Matsuda, M. et al. Recapitulating the human segmentation clock with pluripotent stem cells. Nature 580, 124–129 (2020). 10.1038/s41586-020-2144-9

7 Dequeant, M. L. et al. A complex oscillating network of signaling genes underlies the mouse segmentation clock. Science 314, 1595–1598 (2006). 10.1126/science.1133141

8 Zhang, L., Kendrick, C., Julich, D. & Holley, S. A. Cell cycle progression is required for zebrafish somite morphogenesis but not segmentation clock function. Development 135, 2065–2070 (2008). 10.1242/dev.022673

9 Benazeraf, B. et al. Multi-scale quantification of tissue behavior during amniote embryo axis elongation. Development 144, 4462–4472 (2017). 10.1242/dev.150557

10 Bulusu, V. et al. Spatiotemporal Analysis of a Glycolytic Activity Gradient Linked to Mouse Embryo Mesoderm Development. Dev Cell 40, 331–341 e334 (2017). 10.1016/j.devcel.2017.01.015

11 Mongera, A., Michaut, A., Guillot, C., Xiong, F. & Pourquie, O. Mechanics of Anteroposterior Axis Formation in Vertebrates. Annu Rev Cell Dev Biol 35, 259–283 (2019). https://doi.org/10.1146/annurev-cellbio-100818-125436

12 Diaz-Cuadros, M. et al. Metabolic regulation of species-specific developmental rates. Nature 613, 550–557 (2023). 10.1038/s41586-022-05574-4

13 Carrieri, F. A. et al. CDK1 and CDK2 regulate NICD1 turnover and the periodicity of the segmentation clock. EMBO Rep 20, e46436 (2019). 10.15252/embr.201846436

14 Delaune, E. A., Francois, P., Shih, N. P. & Amacher, S. L. Single-cell-resolution imaging of the impact of Notch signaling and mitosis on segmentation clock dynamics. Dev Cell 23, 995–1005 (2012). 10.1016/j.devcel.2012.09.009

15 Wiedermann, G. et al. A balance of positive and negative regulators determines the pace of the segmentation clock. Elife 4, e05842 (2015). 10.7554/eLife.05842

16 Kicheva, A. et al. Coordination of progenitor specification and growth in mouse and chick spinal cord. Science 345, 1254927 (2014). 10.1126/science.1254927

17 Boehm, B. et al. The role of spatially controlled cell proliferation in limb bud morphogenesis. PLoS Biol 8, e1000420 (2010). 10.1371/journal.pbio.1000420

18 Abe, T. et al. Visualization of cell cycle in mouse embryos with Fucci2 reporter directed by Rosa26 promoter. Development 140, 237–246 (2013). 10.1242/dev.084111

19 El Azhar, Y., et al. Unravelling differential Hes1 dynamics during axis elongation of mouse embryos through single-cell tracking. Development 151 (2024). 10.1242/dev.202936

20 Anderson, M. J. et al. TCreERT2, a transgenic mouse line for temporal control of Cre-mediated recombination in lineages emerging from the primitive streak or tail bud. PLoS ONE 8, e62479 (2013). 10.1371/journal.pone.0062479

21 Sakaue-Sawano, A. et al. Visualizing spatiotemporal dynamics of multicellular cell-cycle progression. Cell 132, 487–498 (2008). 10.1016/j.cell.2007.12.033

22 Palmeirim, I., Henrique, D., Ish-Horowicz, D. & Pourquie, O. Avian hairy gene expression identifies a molecular clock linked to vertebrate segmentation and somitogenesis. Cell 91, 639–648 (1997). 10.1016/s0092-8674(00)80451-1

23 Evrard, Y. A., Lun, Y., Aulehla, A., Gan, L. & Johnson, R. L. lunatic fringe is an essential mediator of somite segmentation and patterning. Nature 394, 377–381 (1998). 10.1038/28632

24 Abe, T. et al. Establishment of conditional reporter mouse lines at ROSA26 locus for live cell imaging. Genesis 49, 579–590 (2011). 10.1002/dvg.20753

25 Yoshioka-Kobayashi, K. et al. Coupling delay controls synchronized oscillation in the segmentation clock. Nature 580, 119–123 (2020). 10.1038/s41586-019-1882-z

26 Pardee, A. B. A restriction point for control of normal animal cell proliferation. Proc Natl Acad Sci U S A 71, 1286–1290 (1974). 10.1073/pnas.71.4.1286

27 Tsiairis, C. D. & Aulehla, A. Self-Organization of Embryonic Genetic Oscillators into Spatiotemporal Wave Patterns. Cell 164, 656–667 (2016). 10.1016/j.cell.2016.01.028

28 Meijer, H. A. et al. NOTCH1 S2513 is critical for the regulation of NICD levels impacting the segmentation clock in hiPSC-derived PSM cells and somitoids. bioRxiv, 2024.2012.2010.627712 (2024). 10.1101/2024.12.10.627712

29 Sonnen, K. F. et al. Modulation of Phase Shift between Wnt and Notch Signaling Oscillations Controls Mesoderm Segmentation. Cell 172, 1079–1090 e1012 (2018). 10.1016/j.cell.2018.01.026

30 Aulehla, A. et al. Wnt3a plays a major role in the segmentation clock controlling somitogenesis. Dev Cell 4, 395–406 (2003). 10.1016/s1534-5807(03)00055-8

31 Greco, T. L. et al. Analysis of the vestigial tail mutation demonstrates that Wnt-3a gene dosage regulates mouse axial development. Genes & A 10, 313–324 (1996).

32 van Oostrom, M. J., Meijer, W. H. M. & Sonnen, K. F. A Microfluidics Approach for the Functional Investigation of Signaling Oscillations Governing Somitogenesis. J Vis Exp 169, 1–17 (2021). 10.3791/62318

33 Masamizu, Y. et al. Real-time imaging of the somite segmentation clock: revelation of unstable oscillators in the individual presomitic mesoderm cells. Proc Natl Acad Sci U S A 103, 1313–1318 (2006). 10.1073/pnas.0508658103

34 Morgan, D. O. The cell cycle : principles of control. (2007).

35 Pourquie, O. The segmentation clock: converting embryonic time into spatial pattern. Science 301, 328–330 (2003). 10.1126/science.1085887

36 Soroldoni, D. et al. Genetic oscillations. A Doppler effect in embryonic pattern formation. Science 345, 222–225 (2014). 10.1126/science.1253089

37 Cooke, J. & Zeeman, E. C. A clock and wavefront model for control of the number of repeated structures during animal morphogenesis. J Theor Biol 58, 455–476 (1976). 10.1016/s0022-5193(76)80131-2

38 Niwa, Y. et al. Different types of oscillations in Notch and Fgf signaling regulate the spatiotemporal periodicity of somitogenesis. Genes Dev 25, 1115–1120 (2011). 10.1101/gad.2035311

39 Dubrulle, J. & Pourquie, O. fgf8 mRNA decay establishes a gradient that couples axial elongation to patterning in the vertebrate embryo. Nature 427, 419–422 (2004). 10.1038/nature02216

40 Dubrulle, J., McGrew, M. J. & Pourquie, O. FGF signaling controls somite boundary position and regulates segmentation clock control of spatiotemporal Hox gene activation. Cell 106, 219–232 (2001). 10.1016/s0092-8674(01)00437-8

41 Mongera, A. et al. A fluid-to-solid jamming transition underlies vertebrate body axis elongation. Nature 561, 401–405 (2018). 10.1038/s41586-018-0479-2

42 Naiche, L. A., Holder, N. & Lewandoski, M. FGF4 and FGF8 comprise the wavefront activity that controls somitogenesis. Proc Natl Acad Sci U S A 108, 4018–4023 (2011). 10.1073/pnas.1007417108

43 Cambray, N. & Wilson, V. Axial progenitors with extensive potency are localised to the mouse chordoneural hinge. Development 129, 4855–4866 (2002). 10.1242/dev.129.20.4855

44 Mathis, L. & Nicolas, J. F. Different clonal dispersion in the rostral and caudal mouse central nervous system. Development 127, 1277–1290 (2000). 10.1242/dev.127.6.1277

45 Neijts, R., Simmini, S., Giuliani, F., van Rooijen, C. & Deschamps, J. Region-specific regulation of posterior axial elongation during vertebrate embryogenesis. Dev Dyn 243, 88–98 (2014). 10.1002/dvdy.24027

46 Yaman, Y. I. & Ramanathan, S. Controlling human organoid symmetry breaking reveals signaling gradients drive segmentation clock waves. Cell 186, 513–527 e519 (2023). 10.1016/j.cell.2022.12.042

47 Thomson, L., Muresan, L. & Steventon, B. The zebrafish presomitic mesoderm elongates through compaction-extension. Cells Dev 168, 203748 (2021). 10.1016/j.cdev.2021.203748

48 Sabherwal, N. et al. Differential phase register of Hes1 oscillations with mitoses underlies cell-cycle heterogeneity in ER(+) breast cancer cells. Proc Natl Acad Sci U S A 118 (2021). 10.1073/pnas.2113527118

49 Weterings, S. D. C. et al. NOTCH-driven oscillations control cell fate decisions during intestinal homeostasis. bioRxiv, 2024.2008.2026.609553 (2024). 10.1101/2024.08.26.609553

50 Maeda, Y., Isomura, A., Masaki, T. & Kageyama, R. Differential cell-cycle control by oscillatory versus sustained Hes1 expression via p21. Cell Rep 42, 112520 (2023). 10.1016/j.celrep.2023.112520

51 Zeng, C. et al. Evaluation of 5-ethynyl-2’-deoxyuridine staining as a sensitive and reliable method for studying cell proliferation in the adult nervous system. Brain Res 1319, 21–32 (2010). 10.1016/j.brainres.2009.12.092

52 Choi, H. M. T. et al. Third-generation in situ hybridization chain reaction: multiplexed, quantitative, sensitive, versatile, robust. Development 145 (2018). 10.1242/dev.165753

