## Supplementary Information for "Coupling of cell proliferation to the segmentation clock ensures robust somite scaling"

Supplementary Information contains 14 Supplementary Figures, Supplementary Theory and 6 Supplementary Movies.

### Supplementary Figure Legends

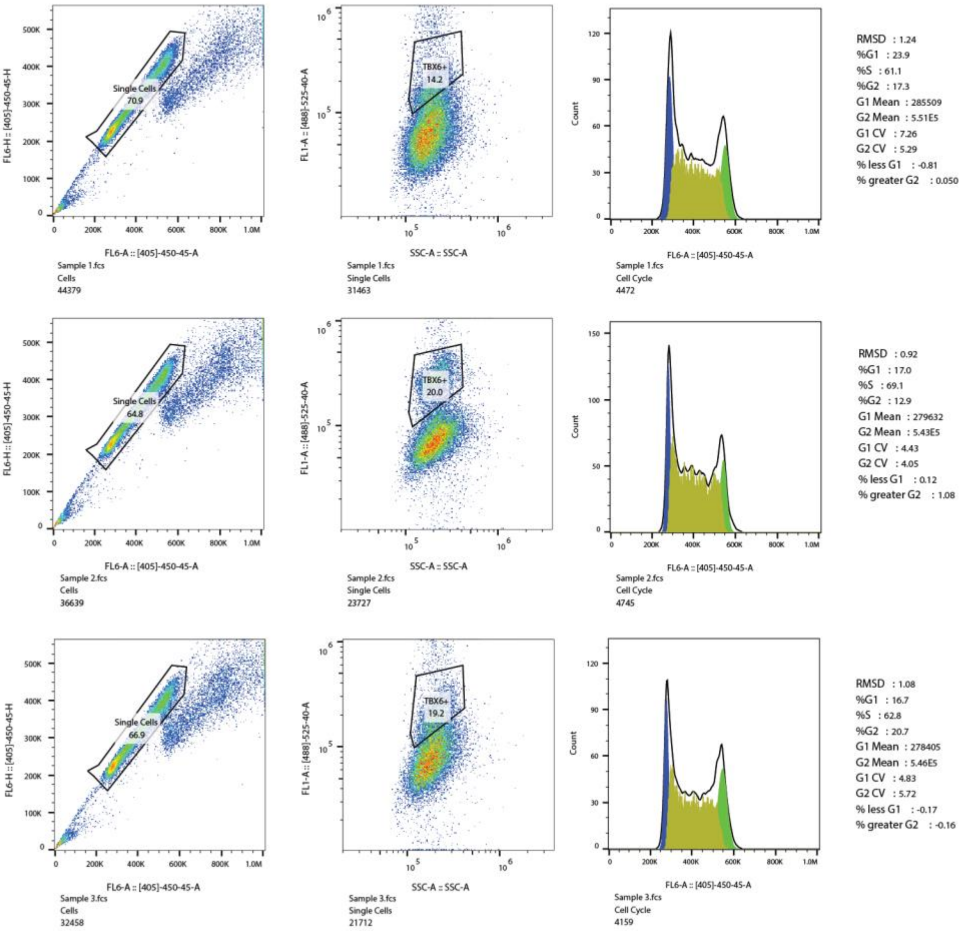


**Figure S1 Quantification of cell cycle phases in PSM cells by flow cytometry.** E10.5 embryonic tails were dissociated into single cells. Immunostaining against Tbx6 and counterstaining with Hoechst were performed before analysis by flow cytometry. Each row represents quantification of one sample. Left panel: Forward- versus side-scatter; gating on single cells. Middle panel: Tbx6 signal versus side-scatter; gating on Tbx6-positive cells. Right panel: histogram of DNA content in Tbx6-positive cells. Automatic detection of G1 phase (dark blue), S phase (olive green) and G2/M phases (light green).

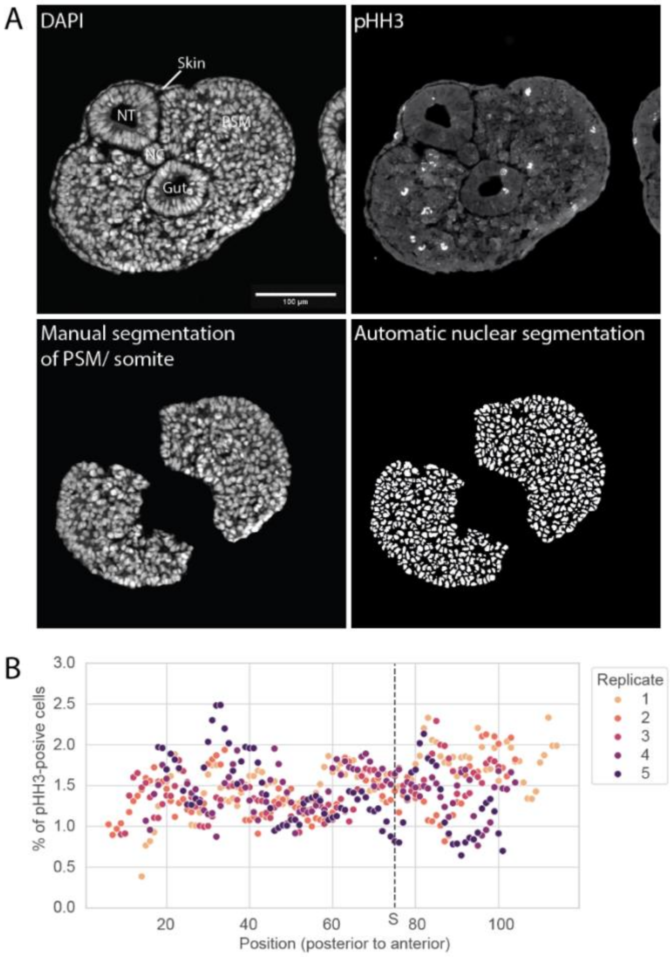


**Figure S2 Quantification of pHH3+ (mitotic) cells in E10.5 presomitic mesoderm.** **A** Representative images of PSM and somitic sections stained for nuclei and PHH3 (as in Figure 1D). Presomitic and somitic regions were manually segmented and nuclei automatically segmented to quantify PHH3-positive cells. **B** Quantification of late mitotic cells in embryonic tails along posterior-anterior axis. N = 5.

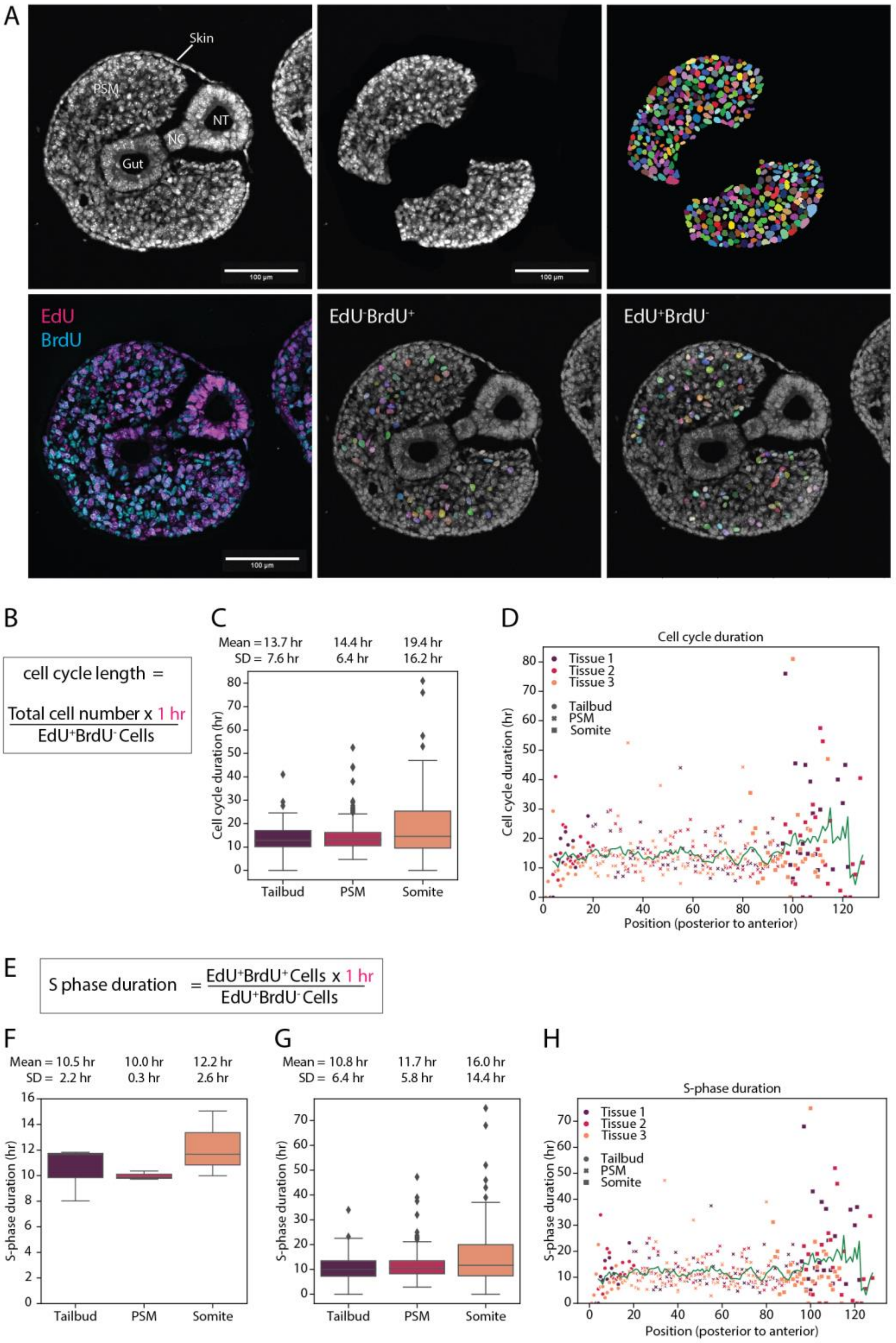


**Figure S3 Quantification of cell cycle dynamics in E10.5 presomitic mesoderm by EdU/BrdU labelling. A** Representative images illustrating the analysis pipeline (as in Figure 1G). Top row: Representative image of stained nuclei in the section (left panel). Presomitic and somitic regions were manually segmented in 5 µm sections stained for nuclei, EdU and BrdU, (middle panel) and nuclei automatically segmented (right panel). Bottom row: Representative image of section stained for EdU and BrdU (image as in Figure 1F). Automatic detection of EdU-BrdU+ (middle panel) and EdU+BrdU- cells (right panel). **B-D** Quantification of cell cycle duration. **B** Formula for the quantification of cell cycle time. **C** Quantification of cell cycle time per section for each region **D** Quantification of cell cycle time along the posterior-anterior axis. Each point corresponds to quantification of individual 5 μm tissue sections. Green line: moving average of 5. **E-H** Quantification of S phase duration. **E** Formula for the quantification of S phase duration. **F** Quantification of S phase duration per region indicated in Figure 1B. **G** Quantification of S phase duration per section in the regions indicated in Figure 1B. **H** Quantification of S phase duration along the posterior-anterior axis. Each point corresponds to quantification of individual 5 μm tissue section. Green line: moving average of 5.

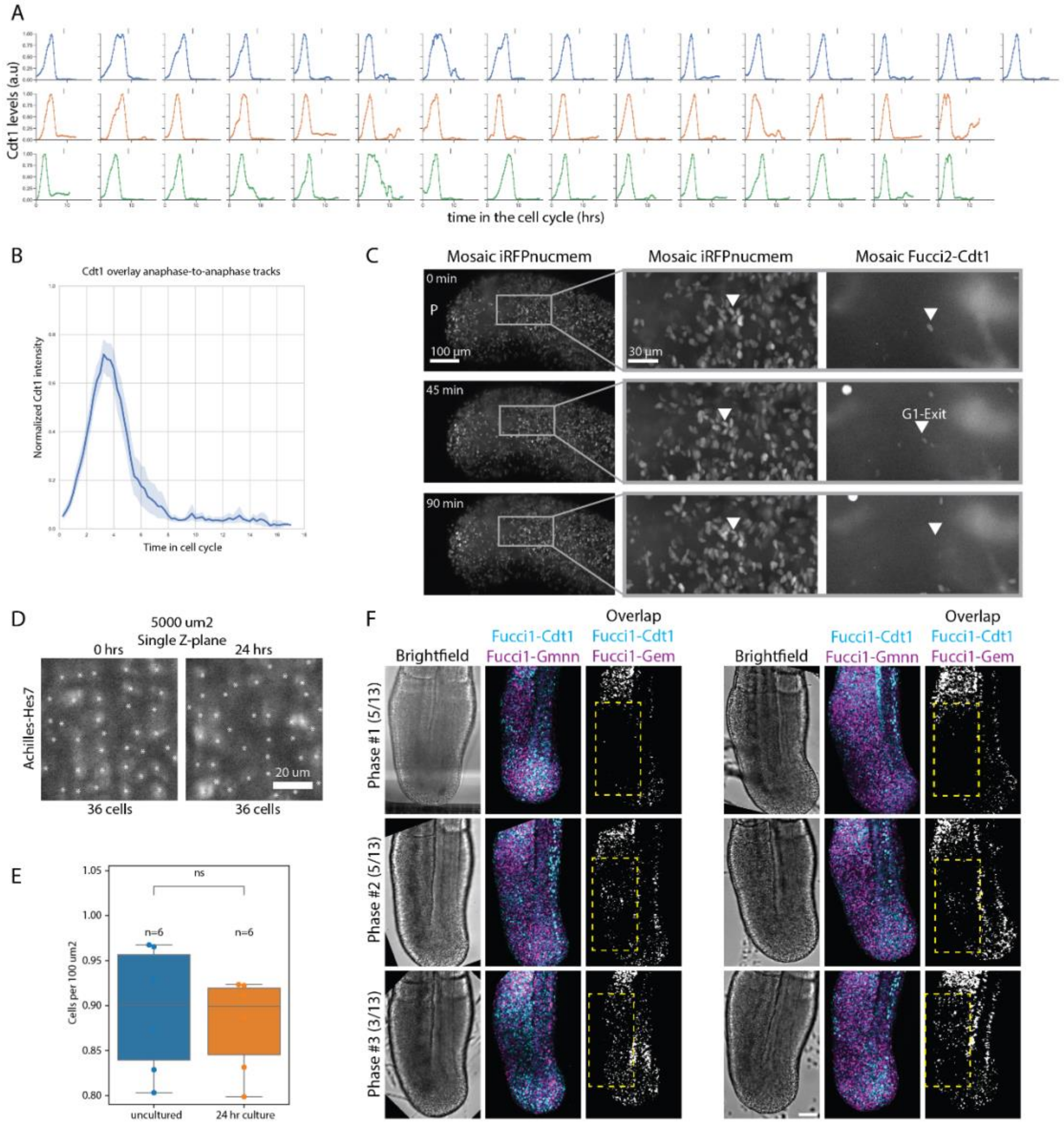


**Figure S4 Cell cycle dynamics in ex vivo cultured E10.5 embryonic tails. A,B** Quantification of Fucci2-Cdt1 signal in tracked single cells (**A**) and as mean (blue line) and standard deviation (light blue shading) (**B**). Corresponds to data shown in Figure 1K-L. **C** Representative snapshots of timeseries data. Arrowhead points at one cell tracked over time. **D,E** Quantification of cell density in *ex vivo* cultured embryo tails**. D** Representative image of a single plane showing nuclear Achilles-Hes7 at two timepoints during the real-time imaging: t=0h (*left panel*), t = 24 h (*right panel*). Scale bar = 20um . **E** Quantification of cell density in anterior PSM in single z planes as shown in D. **F** Patterns of G1-S transition in uncultured E10.5 embryonic tails (partially same as in Figure 1O). Cells double-positive for Fucci1-Cdt1 and Fucci1-Gmnn were determined in the PSM of embryonic tails. Tails were ordered into three phases according to their staining pattern: Phase 1: no double-positive cells in PSM; Phase 2: stripe of double-positive cells in PSM; Phase 3: double-positive cells distributed over PSM. Number in brackets indicates number of tails in which specific staining pattern was observed. 2 sets of representative images are shown for each phase with brightfield (*left panel*), merge of Fucci1-Cdt1 and Fucci1-Gmnn (*middle panel*) and cells double-positive for Fucci1-Cdt1 and Fucci1-Gmnn (*right panel*). N=13. Right set of images corresponds to Fig. 1N. Scale bar 100 μm.

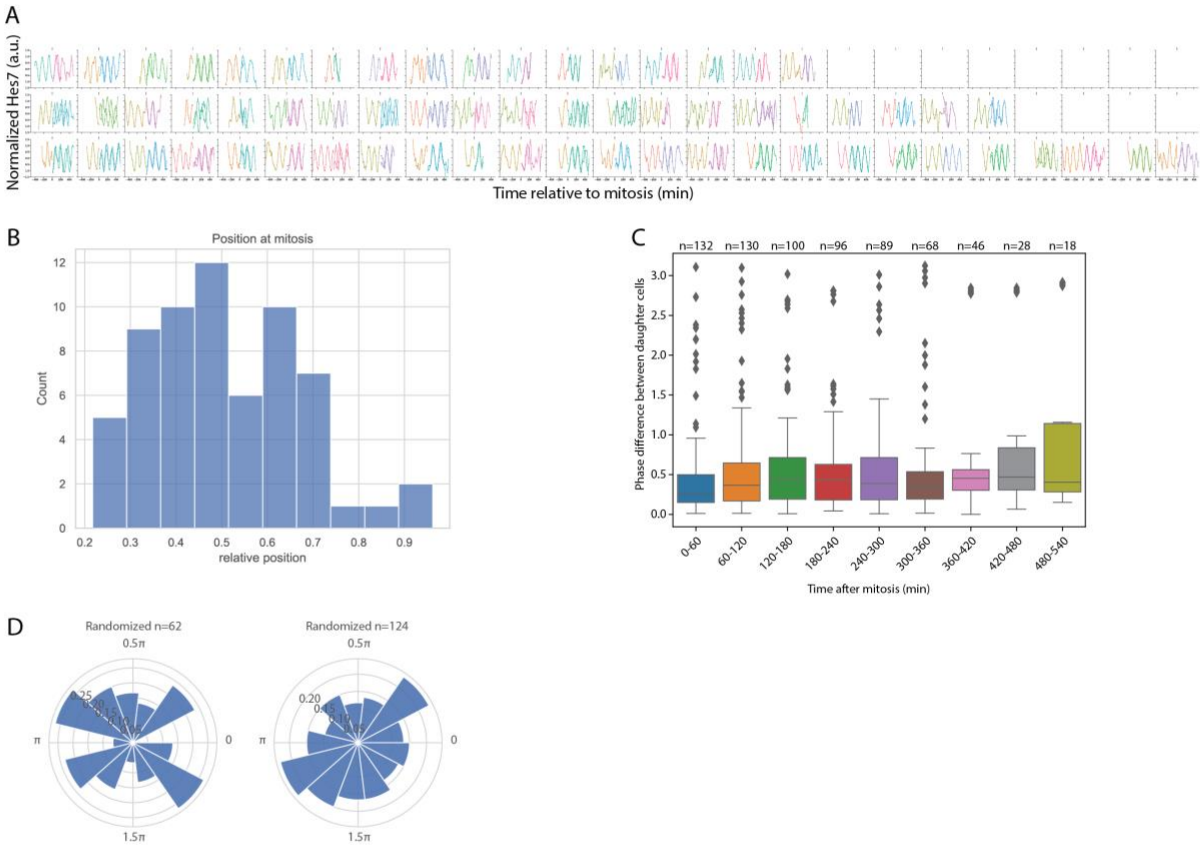


**Figure S5 Quantifying segmentation clock oscillations relative to cell divisions.** E10.5 embryo tails, expressing Achilles-Hes7 and mosaically expressed H2B-mCherry, were cultured *ex vivo* and fluorescence real-time imaging was performed. Single cells were tracked over time. **A** Detrended timeseries data of Achilles-Hes7 expression in tracked single cells. Change in line colour indicates cell division. **B** The length of the PSM is divided into 10 bins from posterior PSM to anterior PSM. Histogram of the positions where mitoses occurred in each of the single-cell tracks. **C** The phase difference of Hes7 oscillations between daughter cells was quantified in dependence of time after mitosis. **D** Representative radial plots showing random phases for 62 (*left panel*) and 124 samples (*right panel*), referring to Fig. 2D.

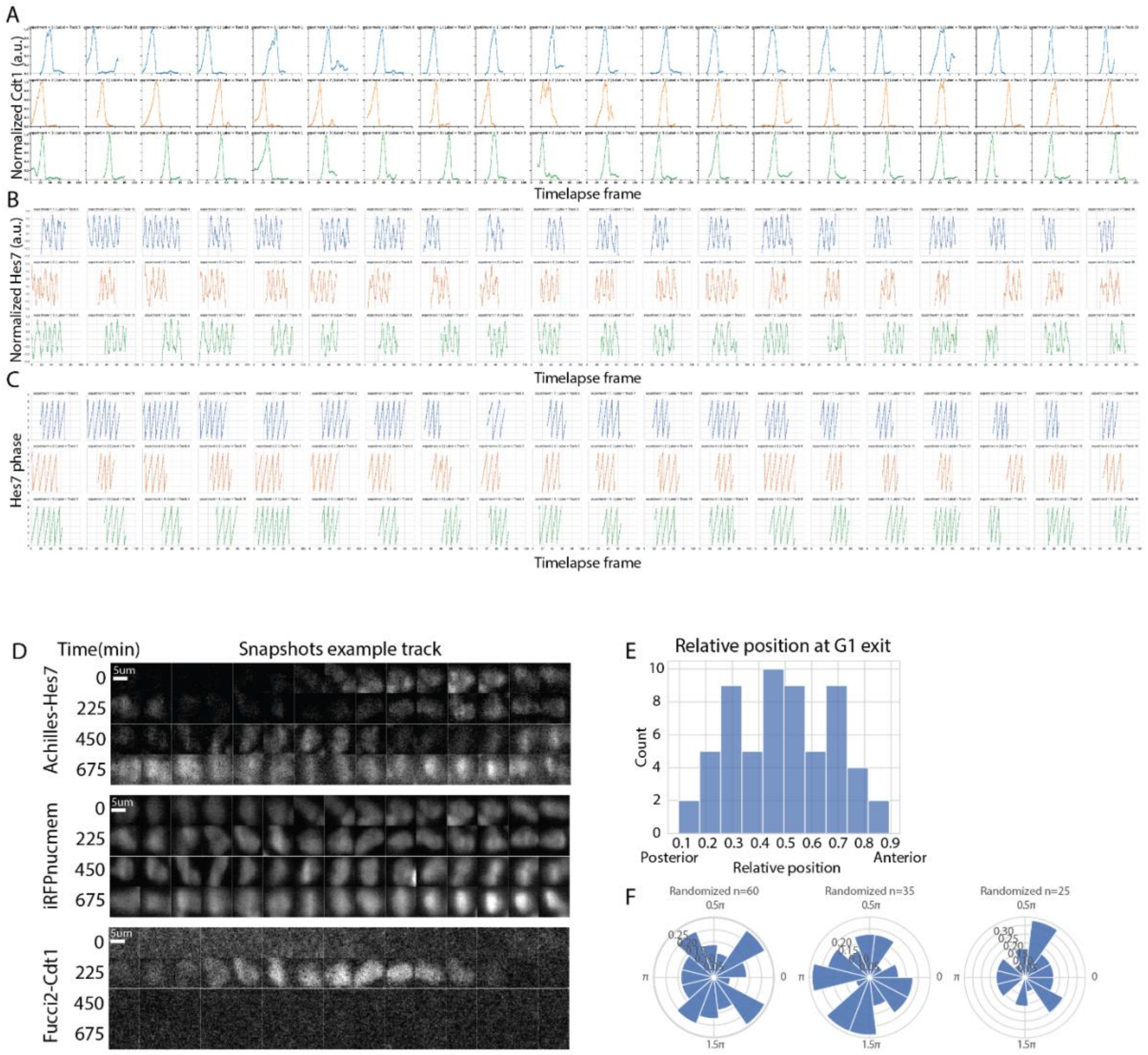


**Figure S6 Quantifying segmentation clock oscillations relative to G1-S transition.** E10.5 embryo tails expressing Achilles-Hes7 and mosaically expressed Fucci2-Cdt1 and iRFPnucmem were cultured *ex vivo* and fluorescence real-time imaging was performed. Single cells were tracked over time. **A** Quantification of Cd1-mCherry signal in tracked single cells. **B** Detrended timeseries data of Achilles-Hes7 expression in tracked single cells. **C** Data for Hes7 oscillations phase for single-cell tracks shown in B. **D** Representative snapshots of timeseries dataset. **E** The length of the PSM is divided into 10 bins from posterior PSM to anterior PSM. Histogram of the positions where G1-S transition occurred in each of the single-cell tracks. **F** Representative radial plots showing random phases for 60 (*left panel*), 35 (*middle panel*) and 25 samples (*right panel*), referring to Fig. 2H.

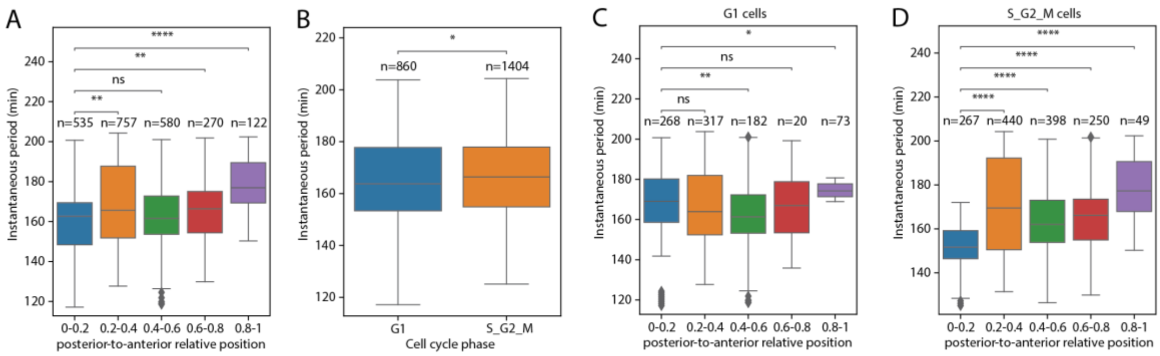


**Figure S7 Analysis of segmentation clock dynamics in single cells along the posterior-to-anterior axis of the PSM.** Further analysis of single cell tracks from Fig. S6. E10.5 embryo tails, expressing Achilles-Hes7 and mosaically expressed Fucci2-Cdt1 and iRFPnucmem, were cultured *ex vivo* and fluorescence real-time imaging was performed. The instantaneous period of Achilles-Hes7 oscillations was quantified. **A** Instantaneous period of all tracked cells relative to position along the posterior-anterior axis of the PSM. **B** Instantaneous period of all tracked cells in G1 phase (Fucci2-Cdt1 positive) or S/G2/M phase (Fucci2-Cdt1 negative) of the cell cycle. **C** Instantaneous period of all tracked cells in G1 phase (Fucci2-Cdt1 positive) in dependence of relative position along the posterior-anterior axis of the PSM. **D** Instantaneous period of all tracked cells in S/G2/M phase (Fucci2-Cdt1 negative) in dependence of relative position along the posterior-anterior axis of the PSM. * < 0.05; ** < 0.01; *** < 0.001; **** <0.0001.

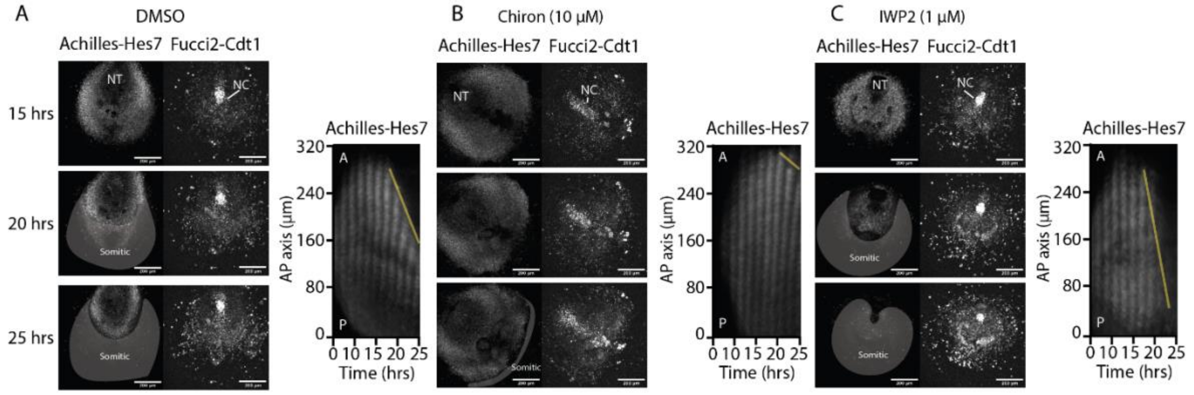


**Figure S8 Modification of Wnt signalling impacts segmentation clock dynamics as expected.** 2D *ex vivo* cultures of E10.5 embryonic tails were incubated with DMSO (**A**), Chiron (**B**) or IWP2 (**C**) and analysed by fluorescence real-time imaging. Representative snapshots of timeseries data are shown in left panels and representative kymographs in right panels. The yellow line indicates regression. Gray shading highlights differentiated somitic cells. While Chiron delays regression IWP2 speeds up regression as seen by the size maintenance or reduction of the oscillating field.

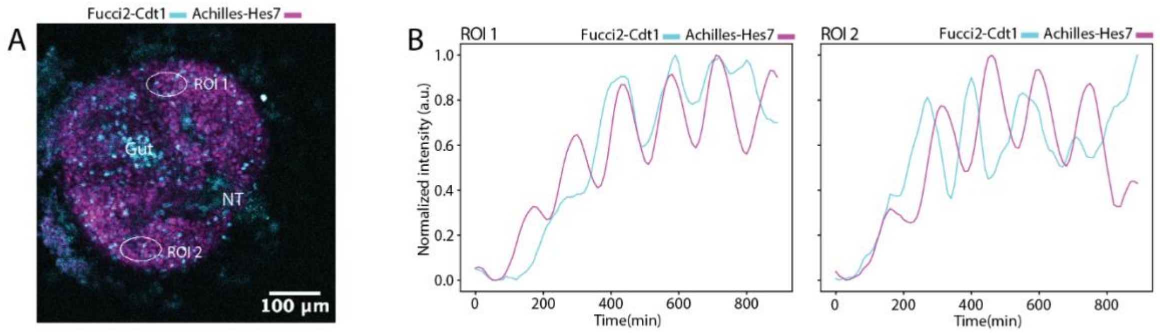


**Figure S9 Correlation of cell cycle dynamics to segmentation clock in 2D *ex vivo* cultures of embryonic tails. A** Maximum Intensity projection of E10.5 embryonic tail (purple: Achilles-Hes7, cyan: Fucci2-Cdt1). **B** Quantification of Achilles-Hes7 and Fucci2-Cdt1 in regions of interest (ROIs) as indicated in A.

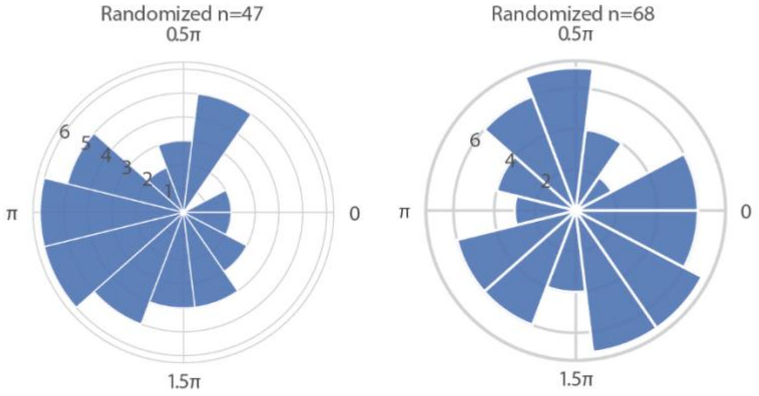


**Figure S10 Quantification of segmentation clock and G1-S transition in single cells within embryonic tails cultured on microfluidic chip.** Representative radial plots showing random phases for 47 (*left panel*) and 68 samples (*right panel*), referring to Fig. 3I.

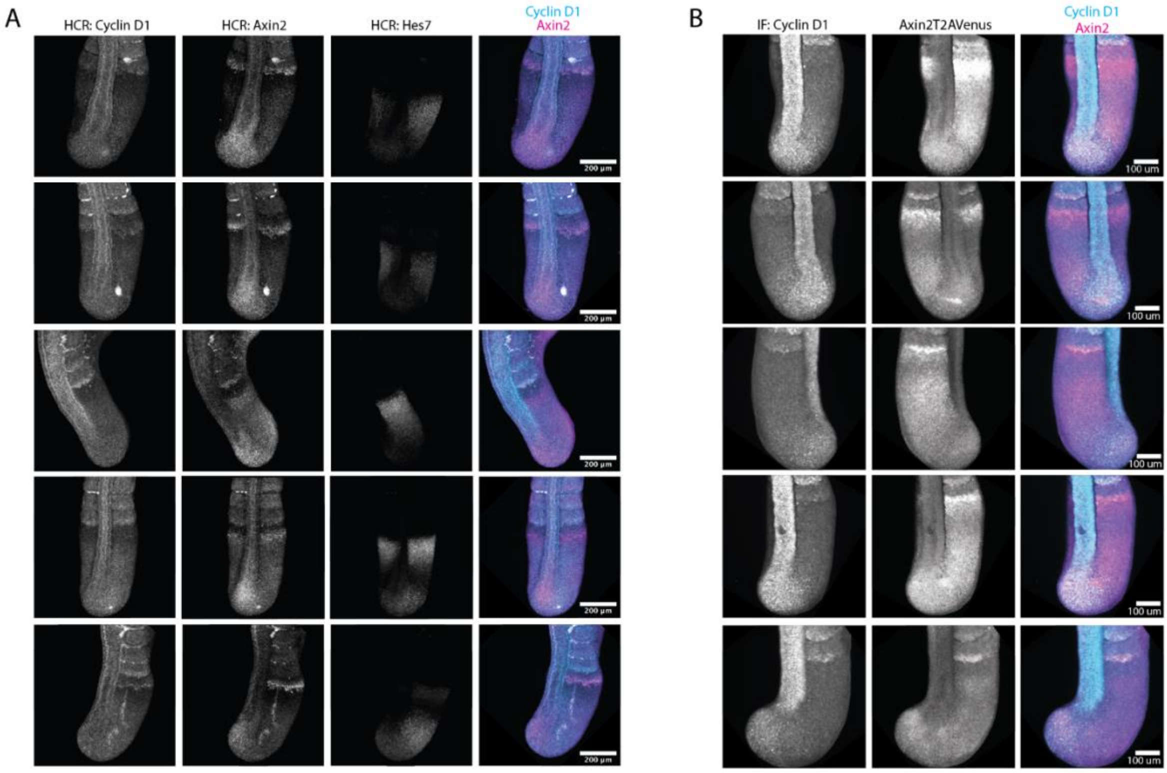


**Figure S11 Changing staining pattern of Cyclin D1 correlates with Axin2 staining.** Representative images are shown. **A** Staining for mRNA levels of Axin2, Cyclin D1 and Hes7 in E10.5 embryonic tails using hybridization chain reaction (HCR) (partially same as in Fig. 3J). **B** Immunostaining against Cyclin D1 in Axin2T2AVenus-expressing E10.5 embryonic tails (partially same as in Fig. 3K).

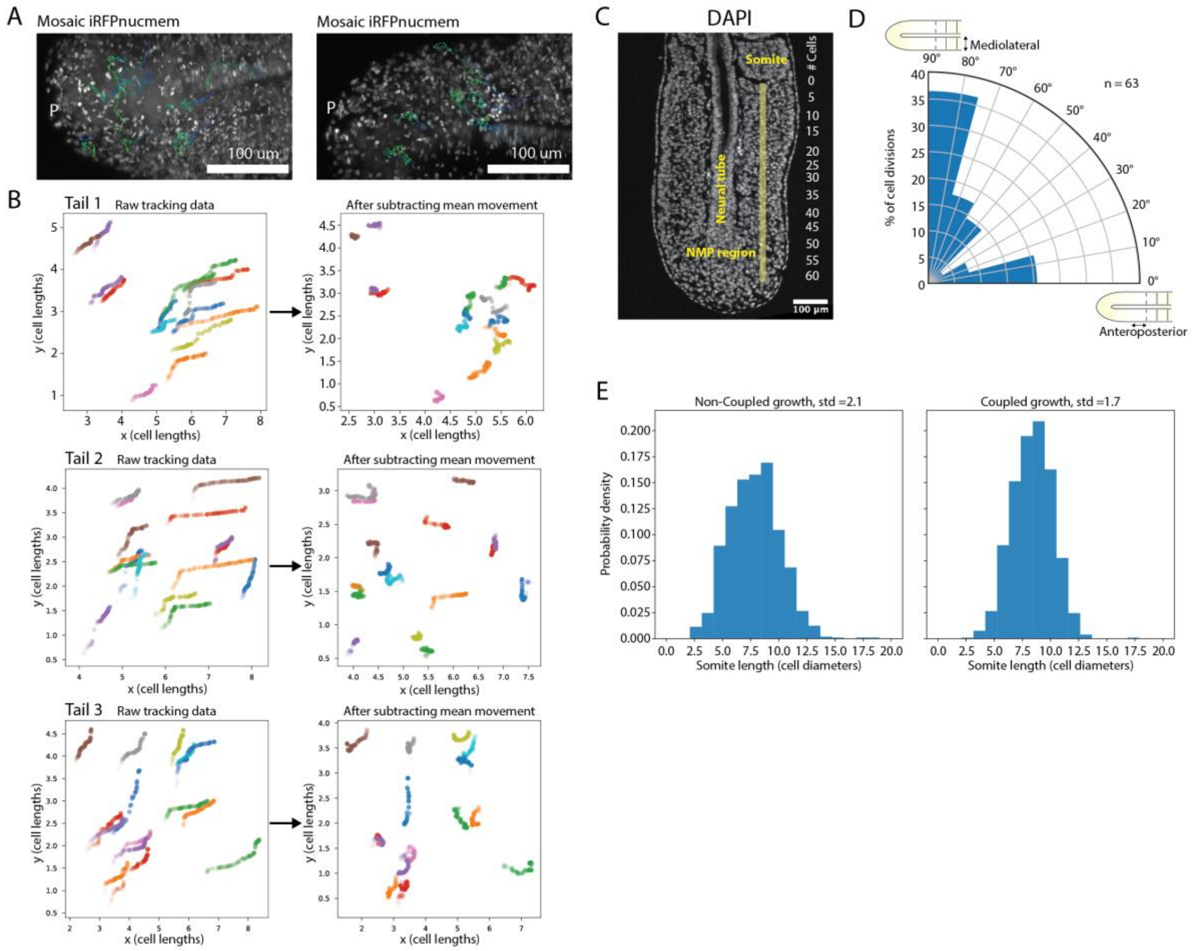


**Figure S12 Quantification of cell motility and PSM length.** E10.5 embryonic tails were cultured ex vivo and fluorescence real-time imaging was performed (corresponds to data from Figure 1,2). **A** Maximum intensity projection of 2 tails with track overlayed onto timepoint 1 of the movie. **B** Plots of single cell tracks from different tails (*left panels*). Whole-tissue movement was computationally removed to reveal movement of single cells relative to their surroundings (*right panels*). N=16 tracks per tail. Tail 1 is the same as in the main figure. **C** Representative 5 µm section of an E10.5 tail counterstained with DAPI. Cells lying on a line (shaded yellow) in an anterior-posterior direction along the PSM were counted. Cell count is given on the right. Cell number from somite boundary to NMP region were considered. **D** Division direction of mitotic cells in Fig. 2A-B relative to the AP axis. **E** Histogram of somite lengths from 100 simulations. std(non-coupled proliferation) = 2.1 cell diameters; std(coupled proliferation) = 1.7 cell diameters.

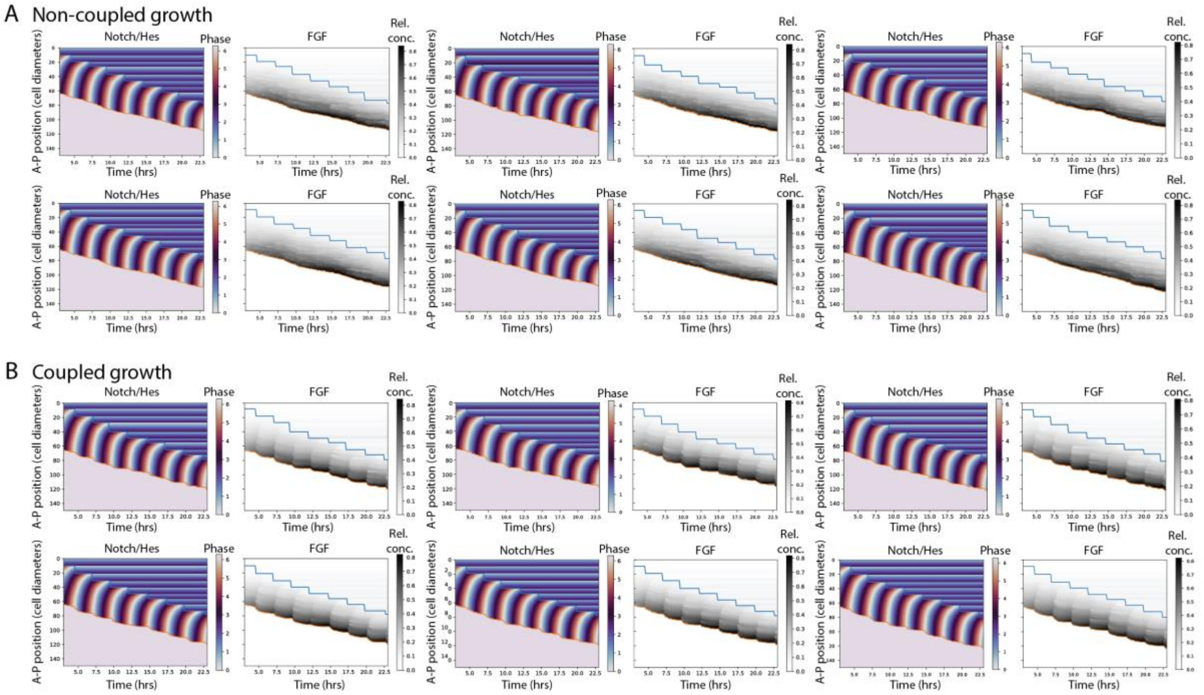


**Figure S13 Simulation of somitogenesis with random cell proliferation and proliferation coupled to the segmentation clock.** Example plots for simulations of signalling waves (*left panels*) as well as FGF gradient and somite formation (*right panels*). **A** Non-coupled cell proliferation. **B** Coupled cell proliferation.

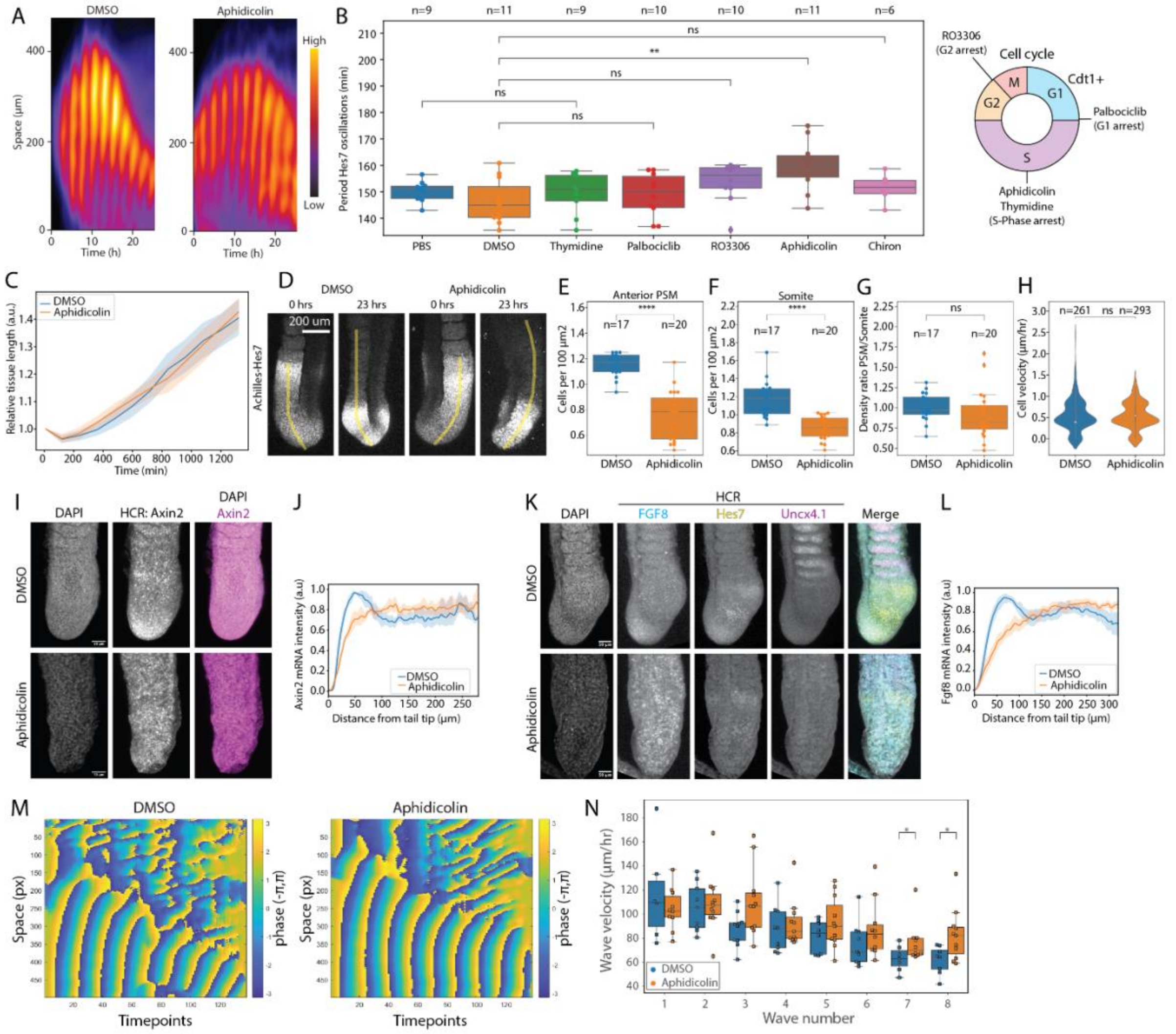


**Figure S14 Effect of cell cycle inhibition on segmentation clock and signalling gradients. A** 2D *ex vivo* cultures of E10.5 embryonic tails were incubated with DMSO (*left panel*) or Aphidicolin (1.6 µg/ml) (*right panel*) and analysed by fluorescence real-time imaging. Representative kymographs are shown. **B** 2D *ex vivo* cultures of E10.5 embryonic tails were incubated with the indicated drugs and analysed by fluorescence real-time imaging. The period of Hes7 oscillations was quantified in anterior PSM. * < 0.05; ** < 0.01; *** < 0.001; **** <0.0001. **C-M** *Ex vivo* cultures of E10.5 embryonic tails were incubated with DMSO (*left panel*) or Aphidicolin (1.6 µg/ml) (*right panel*) for 23 h. **C,D** Quantification of tissue growth by determining tissue length from the posterior tailtip to the last somite boundary present at the start of the experiment and normalizing to the PSM size at the start of the experiment. **D** Representative snapshots of maximum intensity projections showing Achilles-Hes7 in DMSO (*left panels*) and Aphidicolin-treated samples (*right panels*) at early and late timepoints during the experiment. **E-G** Quantification of cell density**. E,F** Quantification of cell density in anterior PSM (**E**) and last formed somites (**F**) in single z planes as shown in D. **G** Ratio between cell density in anterior PSM and somite. **H** Quantification of cell motility in fluorescence real-time imaging data of embryonic tails expressing Achilles-Hes7 (data corresponds to C,D). **I,J** After 23 h of incubation tissue was fixed and HCR against Axin2 and counterstaining with DAPI were performed. **I** Representative images. Scale bar 100 µm. **J** The pattern of Axin2 expression was quantified along the posterior-anterior axis of the PSM and normalized to 0 to 1. n(DMSO) = 4, n(Aphidicolin) = 6. **K,L** After 23 h of incubation tissue was fixed and HCR against Fgf8 and counterstaining with DAPI were performed. **K** Representative images (partially shown in Fig. 5F). Scale bar 100 µM. **L** The pattern of FGF8 expression was quantified along the posterior-anterior axis of the PSM and normalized to 0 to 1. n(DMSO) = 8, n(Aphidicolin) = 13. **M** Phase kymographs of the Achilles-Hes7 fluorescence intensity kymographs shown in Fig. 5I. **N** Quantification of the wave velocity. n(DMSO) = 10, n(Aphidicolin) = 11. * < 0.05.

**Supplementary Movies**

Movie S1. Time-Lapse Imaging of mouse E10.5 tail explant expressing mosaic iRFP-Nucmem (grey) and Fucci2-Cdt1 (cyan) with 15 single cell tracks plotted on top of the movie. Related to Figure 1I-J.

Movie S2. Time-Lapse Imaging of mouse E10.5 tail explant expressing Achilles-Hes7 (magenta) and Fucci2-Cdt1 (cyan). Quantification of the two ROIs can be found in Figure 2K.

Movie S3. Time-Lapse Imaging of mouse E10.5 tail tip culture expressing Achilles-Hes7 (magenta) and Fucci2-Cdt1 (cyan). Quantification of the two ROIs can be found in Figure S9B.

Movie S4. Time-Lapse Imaging of mouse E10.5 tail tip culture during microfluidics experiment. Embryos receive pulses of Chiron (top row) or DMSO (bottom row) when flow is seen in the upper panel. Embryos express Achilles-Hes7 (magenta) and Fucci2-Cdt1 (cyan). Related to Figure 3H-I.

Movie S5. Time-Lapse Imaging of mouse E10.5 tail tip culture exposed to constant cell cycle perturbation. Embryos receive small molecules perturbing progression through specific cell cycle phases. Embryos express Achilles-Hes7 (top row) and Fucci2-Cdt1 (bottom row). Related to Figure S14B. Accumulation of Fucci2-Cdt1 can be observed when blocking with Cdk4/6 inhibitor Palbociclib, while no new appearance of Fucci2-Cdt1 can be observed when blocking with Thymidine, Aphidicolin or RO3306.

Movie S6. Time-Lapse Imaging of mouse E10.5 tail explant perturbed with Aphidicolin (right) or DMSO (left) expressing Achilles-Hes7. Related to Figure 5, S14.
