## Supplementary Theory for "Coupling of cell proliferation to the segmentation clock ensures robust somite scaling"

The understanding of somite formation has evolved significantly over the past century, pioneered by the "Clock and Wavefront" model, proposed by Cooke and Zeeman [1, 2]. The model relies on two key mechanisms: an oscillating genetic "clock" that creates regular pulses of gene expression, and a moving "wavefront" of cellular competence that gradually progresses through the presomitic mesoderm (PSM). The molecular clock involves synchronized cyclic expression of genes of the Wnt, Notch and FGF signaling pathways. The wavefront is established by opposing gradients of FGF/Wnt (highest in posterior) and retinoic acid (highest in anterior). As the wavefront moves posteriorly, it determines where oscillating cells become competent to form somite boundaries. Building on this foundation, Ishimatsu *et al.* [3] proposed the clock and scaled gradient" model to explain the scaling of the somites with PSM size in zebrafish. Further refinements emerged as imaging techniques advanced: Soroldoni *et al.* [4] established the importance of the phase waves of the oscillating genes in creating patterns of increasingly narrower somites, while Yaman *et al* [5] demonstrated that the period of clock oscillations correlates with local FGF density. These advances in our understanding of the clock mechanism raise important questions about its coordination with other cellular processes. In this supplemental section, we will incorporate these elements to present a mathematical model of somitogenesis, with particular focus on how the segmentation clock interacts with the cell cycle - a coupling that we show plays a crucial role in maintaining robust pattern formation.

### T1. MATHEMATICAL MODEL OF SOMITOGENESIS

We propose a mathematical framework that describes the spatiotemporal dynamics of the key molecular players in somitogenesis. The system tracks two primary variables: the phase of segmentation clock oscillations $\theta$ (in phase with Hes and Notch) and the FGF morphogen concentration $\rho$. We represent the PSM as a one-dimensional domain $[0, L]$ in units of cell lengths from the anterior (somite) to the posterior (tail tip) end. As discussed in the main text, there is very little diffusion of the cells so we only model in the anterior-posterior direction where growth and gradients act.

The evolution of the clock phase is given by the following equation,

$$\partial_t \theta + v \partial_x \theta = \omega + \epsilon \partial_x^2 \theta \tag{T1}$$

with Neumann boundary conditions at the anterior and the posterior: $\partial_x \theta(0) = C, \partial_x \theta(L) = 0$, where $C$ is a constant to represent the slowing down of clock oscillations near formed somites. There are three contributions: advection due to tissue growth ($v\partial_x \theta$), local oscillations at frequency $\omega$, and spatial coupling through Delta-Notch signaling ($\epsilon \partial_x^2 \theta$) [6]. The segmentation clock wave emerges from the combination of spatial coupling and the fixed-gradient boundary condition at the anterior end (positive $\partial_x \theta(0)$), as shown in Figure 4(D) and corroborated by a similar model in [7]. While [5] propose an explicit relationship between FGF density and Hes oscillations, we opt for a simpler model of phase wave dynamics due to potential species-specific differences between mice and human cells. This choice is further supported by apparent limitations in their model: it does not account for the period-modifying effects of Delta-Notch coupling between neighboring clocks, and it predicts a two-fold difference in periods between anterior and posterior ends—a prediction that contradicts both their experimental data and ours, which show variations of no more than 20%. Overall, this equation is similar to the one used in [4], with the small distinction that the phase waves in their model stems from a built-in gradient of Hes periods along the PSM that always scales with the PSM size.

The FGF morphogen concentration $\rho$ is governed by the following equation,

$$\partial_t \rho + v \partial_x \rho = D \partial_x^2 \rho - (\kappa + g(\theta))\rho + \alpha(x) \tag{T2}$$

$$\partial_x v = g(\theta) \tag{T3}$$

with Neumann boundary conditions at the anterior and the posterior: $\partial_x \rho(0) = 0, \partial_x \rho(L) = 0$. The first equation incorporates diffusion ($D\partial_x^2 \rho$), advection due to growth ($v \partial_x \rho$), and a combination of space-dependent production ($\alpha(x)$) and two decay terms: baseline degradation ($\kappa$) and dilution due to tissue growth ($g(theta)$). The last equation relates tissue velocity $v$ and the clock-dependent growth rate $g(\theta)$ in the standard way. The FGF gradient arises from a complex interplay of multiple signaling pathways, including Wnt and retinoic acid (RA), which are not explicitly represented in our model. To maintain parsimony while capturing the shape of the posterior-to-anterior gradient, we implement a spatially-dependent production rate $\alpha(x)$ with elevated production at the posterior: $\alpha(x) = \alpha_0 \left(\exp(-(1 - x/L)/l_0) - \exp(-1/l_0)\right) / (1 - \exp(-1/l_0))$, parameterized by the maximum production rate $\alpha_0$ and the dimensionless length $l_0$. This functional form ensures a monotonic gradient consistent with experimental observations while avoiding additional mechanistic assumptions about pathway interactions.

As demonstrated in Fig 3(C) of the main text, mitosis exhibits a $\pi$ phase shift relative to the segmentation clock, consistent with observations by [8] that cell division occurs during low Hes expression. Importantly, we focus on mitosis timing for tissue growth calculations rather than the G1-S transition, which is phase-locked to Wnt oscillation and phase-shifted from Hes/Notch oscillations [9]. Furthermore, we take into account the effect of the refractory period of the cell cycle by blocking re-entry into cell cycle immediately after mitosis. The exact implementation will be described in the next paragraph. While cells are allowed to divide, we model the Hes-dependent cell division with $g(\theta)$ as a Gaussian function centered at $\pi$ with a width $\sigma$ that determines the width of the phase window in which cells are more likely to divide,

$$g(\theta) = \Omega_0(2\pi\sigma)^{-1/2}\exp\left[-(\theta-\pi)^2/(2\sigma^2)\right] \tag{T4}$$

where $\Omega_0 = 2\pi/T_{G1}$ with $T_{G1}$ the duration of the G1 phase.

To incorporate statistical fluctuations, we discretize the system such that each grid point represents an ensemble of $N$ cells. As demonstrated in Figure 4(B) of the main text, the system exhibits a remarkably low cell diffusion constant of $D = 0.0014\,(\text{cell lengths})^2/\text{hr}$, indicating minimal mixing between adjacent cell populations along the PSM axis. On the other hand, we assume that the specification of segmentation likely involves collective behaviors of more than one cell. So, we assume that a position is competent to differentiate when the mean FGF density across a local group of $N$ cells falls below a critical value. For each of the $N$ cells in a gridpoint, we track the time since the last division individually. Once the refractory period passes for a cell, it is allowed to divide with a probability rate $g(\theta)$. Then we collect all the division events for those N cells to calculate the mean expansion rate of the tissue.

Finally it is important to note that the model is not meant to be a complete description of somitogenesis, but rather a starting point for further investigation. It is designed to capture the essential features of the segmentation clock and its interaction with the cell cycle, and to provide a framework for understanding the complex dynamics of somitogenesis.

### T2. PARAMETER VALUES

The parameters in the simulations are summarized in the table below

| **Parameter** | **Value** |
|---|---|
| Angular frequency of the clock oscillations $\omega$ | $2\pi/2.5$ hr$^{-1}$ |
| Interaction strength of the clock oscillations $\epsilon$ | 40 (cell lengths)$^2$/hr |
| Boundary condition $C$ at anterior | 0.3 (cell lengths)$^{-1}$ |
| Diffusion constant of cells $D$ | 0.0014 (cell lengths)$^2$/hr |
| Degradation rate of FGF $\kappa$ | 0.2 hr$^{-1}$ |
| Production rate of FGF in tail tip $\alpha_0$ | 0.2 hr$^{-1}$ |
| Dimensionless length of FGF gradient $l_0$ | 0.65 |
| Length of G1 phase $T_{G1}$ | 4.4 hrs |
| Length of S-G2-M phase $T_{S-G2-M}$ | 9.2 ± 0.5 hrs |
| Width of the Hes phase where cell divisions can occur $\sigma$ | 0.1 |
| Number of cells per grid point $N$ | 5 cells |

TABLE I. Model parameters used in numerical simulations for Fig 4(D-E).

$\omega, T_{G1}, T_{S-G2-M}$ are obtained directly from experimental measurements, as explained in the main text and Figure 1. The diffusion constant comes from experimental measurements of cell movements in the PSM, as shown in Figure 4(B). $\sigma$ is estimated to be small as cell divisions are strongly phase locked with Hes low (increasing $\sigma$ does not qualitatively change the results of the simulations). The interaction strength $\epsilon$ and gradient $C$ are fitted by qualitative likeliness to the live-image kymograph of Hes oscillations.

The remaining parameters are estimated via parameter scanning of the model near the biologically relevant regime. We observed that the FGF degradation rate is approximately of the same order of magnitude as the cell cycle frequency, although the exact value is difficult to measure. The production rate must approximately match the degradation rate to maintain homeostasis, thus for simplicity, we set the production rate $\alpha_0$ and the degradation rate $\kappa$ to be equal. The dimensionless length of the FGF gradient is estimated from the FGF gradient length measured in experiments to be of order 1 (see Figure 5(G)). We explore these two degree of freedoms in parameter space by running 30 simulations for each set of parameters and calculating the mean ratio of the somite length to the PSM length as well as the

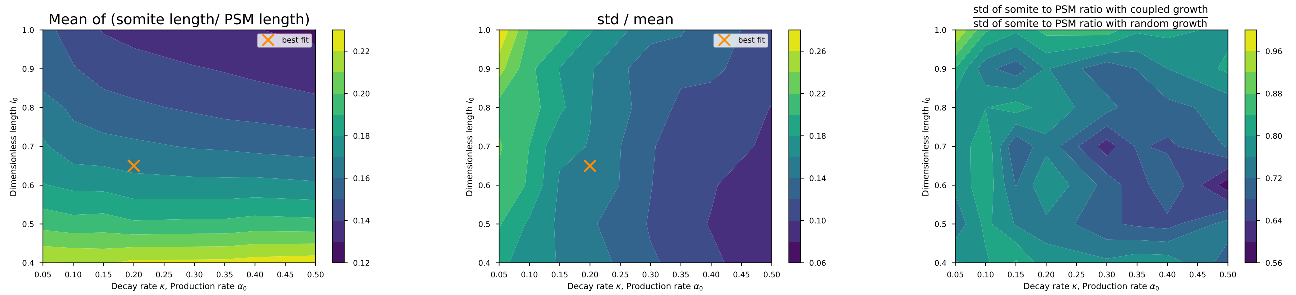


(a) Somite length / PSM length (b) Somite length / PSM length (c) Robustness across parameters

FIG. T1. Parameter scanning in the biologically relevant regime. (a, b) Fitting the mean and standard deviation of the somite length / PSM length ratio to the experimental data. (c) Comparisons between coupled growth and random growth in terms of the standard deviation of the somite to PSM ratio distribution. For all three panels, the rest of the parameters are the same as in Table I and 30 simulations were run for each parameter combination.

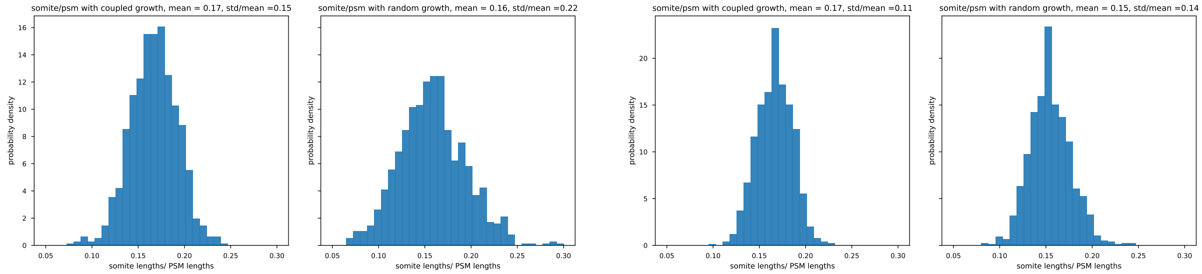


(a) Somite/PSM ratio for 5 cells per grid point (b) Somite/PSM ratio for 10 cells per grid point

FIG. T2. The distributions of (somite length/PSM length) for 5 cells and 10 cells per grid point. The distributions are obtained by collecting data over 100 simulations. The rest of the parameters are the same as in Table I.

standard deviation of the ratio distribution, as shown in Figure T1a and T1b. In experiments, the mean somite to PSM ratio is measured to be approximately 16% - 17 % and the standard deviation / mean is approximately 15% as shown in Figure 5(E). We choose $l_0 = 0.65$ and $\kappa = \alpha_0 = 0.2$ hr$^{-1}$ to match quantities. Furthermore, even without matching the mean and standard deviation to experiments, the coupled growth model consistently yields a tighter distribution of somite lengths compared to random growth across the entire regime of parameters explored (see Figure T1c), implying that the increase in regularity from the coupled growth model is robust to changes in parameters. Finally, we estimate $N$ to be a small number as cell-cell communications across long distance is unlikely. As shown in Figure T2a and T2b, coupled growth provides an improvement in the regularity of the somite length distribution for both values of $N$. Meanwhile we note that the amount of improvement compared to the random growth model depends on the amount of stochasticity in the system: as $N$ – the number of cells per grid point – increases, the gap between the two growth model becomes smaller, as shown in Figure T2a and T2b.

We refer to the main text for further simulation results at the best-fit parameters and discussions.

### T3. PERTURBATION

To mimic the perturbation experiment where we switch off cell cycle, we set the angular frequency of the cell cycle $\Omega_0$ to be 0. The rest of the parameters are the same as in Table I. Note that since all the noise previously described comes from cell divisions, the perturbed case is deterministic and hence we will not discuss noise in this case. Furthermore, since the cells elongate from cell migration even in the absence of cell divisions, we will only be concerned with the ratio of the somite length to the PSM length instead of absolute measurements. With these limitations in mind, we refer back to the main text for further discussions of the simulation results in the no-growth scenario.

---


[1] J. Cooke and E. C. Zeeman, "A clock and wavefront model for control of the number of repeated structures during animal morphogenesis," *Journal of theoretical biology*, vol. 58, no. 2, pp. 455–476, 1976.

[2] O. Pourquié, "The segmentation clock: converting embryonic time into spatial pattern," *Science*, vol. 301, no. 5631, pp. 328–330, 2003.

[3] K. Ishimatsu, T. W. Hiscock, Z. M. Collins, D. W. K. Sari, K. Lischer, D. L. Richmond, Y. Bessho, T. Matsui, and S. G. Megason, "Size-reduced embryos reveal a gradient scaling-based mechanism for zebrafish somite formation," *Development*, vol. 145, no. 11, p. dev161257, 2018.

[4] D. Soroldoni, D. J. Jörg, L. G. Morelli, D. L. Richmond, J. Schindelin, F. Jülicher, and A. C. Oates, "A doppler effect in embryonic pattern formation," *Science*, vol. 345, no. 6193, pp. 222–225, 2014.

[5] Y. I. Yaman and S. Ramanathan, "Controlling human organoid symmetry breaking reveals signaling gradients drive segmentation clock waves," *Cell*, vol. 186, no. 3, pp. 513–527, 2023.

[6] K. Yoshioka-Kobayashi, M. Matsumiya, Y. Niino, A. Isomura, H. Kori, A. Miyawaki, and R. Kageyama, "Coupling delay controls synchronized oscillation in the segmentation clock," *Nature*, vol. 580, no. 7801, pp. 119–123, 2020.

[7] T. Sato, Y. I. Li, D. J. Jorg, M. Komeya, H. Yamanaka, H. Nakamura, K. Hirano, Y. Kondo, K. Aoki, M. Takahashi, *et al.*, "Self-organization of spermatogenic wave coordinates sustained sperm production in the mouse testis," *bioRxiv*.

[8] E. A. Delaune, P. François, N. P. Shih, and S. L. Amacher, "Single-cell-resolution imaging of the impact of notch signaling and mitosis on segmentation clock dynamics," *Developmental cell*, vol. 23, no. 5, pp. 995–1005, 2012.

[9] K. F. Sonnen, V. M. Lauschke, J. Uraji, H. J. Falk, Y. Petersen, M. C. Funk, M. Beaupeux, P. François, C. A. Merten, and A. Aulehla, "Modulation of phase shift between wnt and notch signaling oscillations controls mesoderm segmentation," *Cell*, vol. 172, no. 5, pp. 1079–1090, 2018.
